# Quantitative single-molecule three-color Förster resonance energy transfer by photon distribution analysis

**DOI:** 10.1101/372730

**Authors:** Anders Barth, Lena Voith von Voithenberg, Don C. Lamb

## Abstract

Single-molecule Förster resonance energy transfer (FRET) is a powerful tool to study conformational dynamics of biomolecules. Using solution-based single-pair FRET by burst analysis, conformational heterogeneities and fluctuations of fluorescently labeled proteins or nucleic acids can be studied by monitoring a single distance at a time. Three-color FRET is sensitive to three distances simultaneously and can thus elucidate complex coordinated motions within single molecules. While three-color FRET has been applied on the single-molecule level before, a detailed quantitative description of the obtained FRET efficiency distributions is still missing. Direct interpretation of three-color FRET data is additionally complicated by an increased shot noise contribution when converting photon counts to FRET efficiencies. However, to address the question of coordinated motion, it is of special interest to extract information about the underlying distance heterogeneity, which is not easily extracted from the FRET efficiency histograms directly. Here, we present three-color photon distribution analysis (3C-PDA), a method to extract distributions of inter-dye distances from three-color FRET measurements. We present a model for diffusion-based three-color FRET experiments and apply Bayesian inference to extract information about the physically relevant distance heterogeneity in the sample. The approach is verified using simulated data sets and experimentally applied to triple-labeled DNA duplexes. Finally, 3C-FRET experiments on the Hsp70 chaperone BiP reveal conformational coordinated changes between individual domains. The possibility to address the co-occurrence of intramolecular distances makes 3C-PDA a powerful method to study the coordination of domain motions within biomolecules during conformational changes.

**Significance:** In solution-based single-molecule Förster resonance energy transfer (FRET) experiments, biomolecules are studied as they freely diffuse through the observation volume of a confocal microscope, resulting in bursts of fluorescence from single molecules. Using three fluorescent labels, one can concurrently measure three distances in a single molecule but the experimentally limited number of photons is not sufficient for a straight-forward analysis. Here, we present a probabilistic framework, called three-color photon distribution analysis (3C-PDA), to extract quantitative information from single-molecule three-color FRET experiments. By extracting distributions of interdye distances from the data, the method provides a three-dimensional description of the conformational space of biomolecules, enabling the detection of coordinated movements during conformational changes.

## Introduction

Förster resonance energy transfer (FRET) is a powerful tool to measure intra- or intermolecular distances in the range of 2-10 nm on the single-molecule level. It has been widely applied on the single-molecule level to study the conformational landscape of proteins and nucleic acids using surface-immobilization (1–4) or in solution (5–9). In the latter case, molecules are studied at picomolar concentrations as they diffuse through the observation volume of a confocal microscope, resulting in bursts of fluorescence signal from single molecules (Figure 1A-B) (10–13). The photons from a single molecule burst are analyzed and the molecule-wise distribution of FRET efficiencies provides information regarding the conformational states and dynamics of the biomolecules. Extensions of this method have included lifetime and polarization information (9) as well as direct probing of the acceptor fluorophore using alternating excitation schemes (14, 15) to increase sorting capabilities and informational content of the measurements. Accurate FRET measurements require carefully determined correction factors for spectral crosstalk, direct excitation of the acceptor fluorophore, and differing photon detection efficiencies and quantum yields (16). Using the methods of alternating laser excitation (ALEX) or pulsed interleaved excitation (PIE), these correction factors can be determined from the experiment directly (14, 17). Diffusion-based FRET measurements avoid possible artifacts related to surface immobilization. This advantage comes at the cost of a limited detection time of a few milliseconds per molecule, restricted by the diffusion time through the observation volume as well as fewer photons and thus increased shot noise. The shot-noise contribution can be accounted for in the analysis. Thus, it is possible to elucidate the contributions of physically relevant broadening of the FRET efficiency histogram (18, 19) to study static conformational heterogeneity as well as dynamical interconversion between distinct states (20, 21). This method, named the photon distribution analysis (PDA), has been widely applied in the field (22–29).

**Figure 1:**
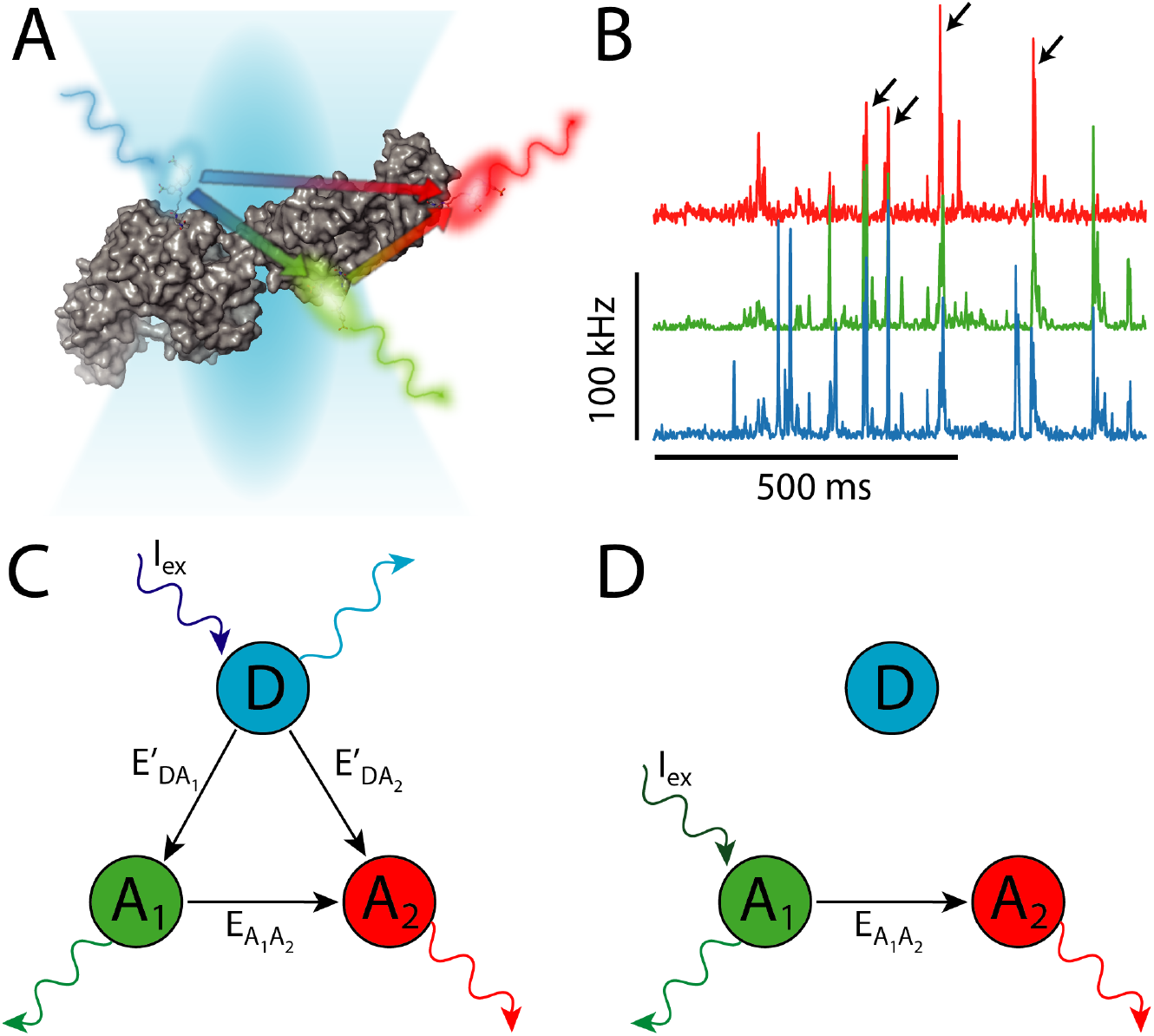
Single-molecule three-color FRET. (A) In a solution-based three-color FRET experiment, single-molecules are measured as they diffuse through the observation volume of a confocal microscope. Possible transition pathways in the three-color FRET system after excitation of the blue dye are schematically shown on a two-domain protein (DnaK, PDB: 2KHO). (B) Single-molecule events result in bursts of fluorescence. Shown are the fluorescence time traces of the combined signal after excitation of the blue, green and red dyes, which are alternatingly excited using pulsed-interleaved excitation (PIE, see Figure S2). Coinciding bursts in all three channels, belonging to triple labeled molecules, are indicated by arrows. (C) Scheme of the possible transition pathways in a three-color FRET system following excitation of the donor dye (blue). Energy can be transferred to either acceptor A_1_ or A_2_ (green and red, respectively) with transition probabilities E’_DA_1__ and E’_DA_2__. Acceptor A1 may further transfer the energy to A_2_ with the FRET efficiency E_A_1_A_2__. (D) Possible transition pathways following direct excitation of acceptor A_1_. Excitation of A_1_ reduces the complexity of the system to independently determine the FRET efficiency between the two acceptors E_A_1_A_2__. Experimentally, blue and green excitation are alternated on the nanosecond timescale using pulsed interleaved excitation (PIE, see Figure S2).

Two-color FRET is limited to the observation of a single distance at a time. Multiple distance readouts are thus only available in separate experiments. Three-color FRET enables the monitoring of three distances in one experiment (30–34). While the same distance distributions are obtainable from three two-color FRET experiments, information about the correlation of distance changes and thus the coordination of molecular movements is only obtained using three-color FRET. This is possible since three-color FRET experiments contain information about the co-occurrence of distances for the individual FRET sensors. During the last decade, a number of single-molecule multicolor FRET studies have been performed both in solution and using surface immobilization (35–41). Proof-of-principle experiments showing the possibility to extract accurate distances from solution-based experiments, however, have been limited to the extraction of average values (35, 36), while a complete description of the shape of the FRET efficiency histogram is still missing. Three-color FRET using surface immobilization has been applied to study coordinated motion in different nucleic acid systems (36–39) and the oligomerization state of proteins (42). With the development of new strategies for protein labeling (43–46), it is becoming more feasible to site-specifically label proteins with three or more fluorophores, opening up the possibility to study coordinated movements within single proteins during their conformational cycle.

However, the extension of two-color FRET to three-color FRET is not as straightforward as it appears and there are many challenges that limit the quantitative analysis of three-color FRET data. First of all, the signal fractions in a three-color FRET experiment are not easily interpreted in terms of distances, as is the case in a two-color FRET experiment. Moreover, any heterogeneity observed for these quantities is difficult to relate to physically relevant distance heterogeneity of the studied system. To complicate the issue, shot noise is a larger problem in three-color FRET measurements because signal after excitation of the donor dye is distributed among three channels, lowering the achievable signal-to-noise ratio. Fluorophores, detectors and optics have been optimized for the visual region of the spectra, which minimizes the obtainable spectral separation of the three channels used for three-color FRET. Hence, larger corrections need to be applied to account for spectral crosstalk and direct excitation of the fluorophores. Thus, extracting accurate distances from three-color FRET experiments remains challenging. To this end, a detailed statistical analysis of the experiment with respect to the underlying three-dimensional distance distribution is needed, which opens up the possibility to study coordinated movements within single biomolecules by analyzing correlations between measured distances.

Here, we present three-color photon distribution analysis (3C-PDA), an extension of the framework of photon distribution analysis to three-color FRET systems. The method is based on a likelihood function for the three-color FRET process in diffusion-based experiments. We apply model-based Bayesian inference using the Metropolis-Hastings algorithm (47, 48) to infer the model parameters. The method is tested using synthetic data and applied experimentally to triple-labeled doublestranded DNA as a model system and to investigate the coordinated motions with the Hsp70 chaperone BiP.

## Theory

### Three-color FRET

FRET is the non-radiative energy transfer from a donor fluorophore *D* to an acceptor fluorophore *A* mediated by dipole-dipole interactions with a strong dependence on distance (49):

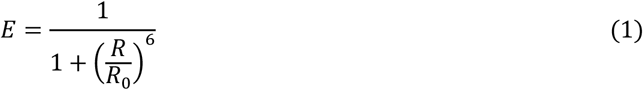

where *R* is the interdye distance and *R*_0_ is the Förster distance which depends on the quantum yield of the donor fluorophore Φ_*D*_, the spectral overlap integral *J*, the orientation of the donor and acceptor fluorophore through the factor *κ*^2^ and the refractive index of the surrounding medium *n*:

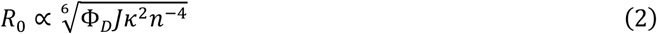

In the experiment, the amount of energy transfer can be measured by separating the fluorescence signal according to the emission spectra of the donor and acceptor fluorophore. A qualitative measure of the FRET efficiency can be obtained from the raw signal fractions in the donor and acceptor channels after donor excitation 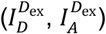, which we call the proximity ratio *PR*:

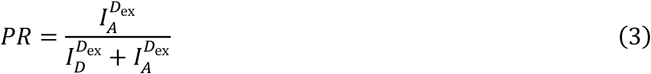

When alternating laser excitation is employed using PIE or ALEX (17, 50), accurate intensity-based FRET efficiencies can be determined when appropriate corrections are applied to the photon counts (17):

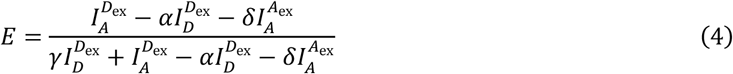

where α is the correction factor for spectral crosstalk of the donor in the acceptor channel, δ is the correction factor for direct excitation of the acceptor at the wavelength of the donor, γ is the correction factor for different quantum yields and detection efficiencies of the fluorophores, and 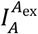 is the signal in the acceptor channel after acceptor excitation.

In three-color FRET, energy can be transferred from a donor fluorophore *D* to two spectrally different acceptor fluorophores *A*_1_ and *A*_2_ (see Figure 1C-D). One of the acceptors, which we define as *A*_1_, may also transfer energy to the lower energy acceptor *A*_2_. Thus, the signal observed for *A*_2_ contains contributions from two possible pathways, either from direct energy transfer from *D*, or from two-step energy transfer mediated by *A*_1_. The signal fractions after excitation of the donor dye are defined by:

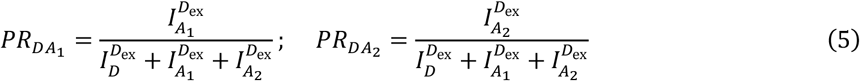

It should be noted that, contrary to two-color FRET, these quantities are not directly related to distances.

The FRET efficiency is defined as the fraction of transitions from the excited donor dye to the acceptor dye, given by:

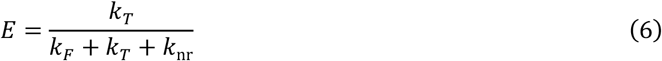

where *k_T_* is the rate of energy transfer, *k_F_* is the intrinsic radiative decay rate of the donor, and *k*_nr_ is the sum over all non-radiative relaxation pathways of the donor. In three-color FRET systems, energy transfer may occur from the donor dye to either acceptor. The efficiency of energy transfer to one acceptor dye is thus modified by the quenching effect of the second acceptor dye, effectively reducing the Förster radius of the dye pair. In analogy to equation 6, one can define apparent FRET efficiencies from the donor dye to either acceptor 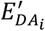, which describe the transition probabilities in the three-color FRET system as shown in Figure 1C.

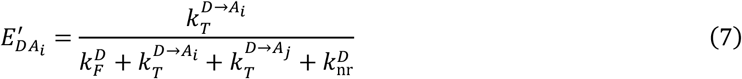

The total FRET efficiency from the donor dye to both acceptor dyes is then given by:

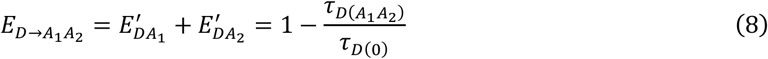

where *τ*_*D*(0)_ is donor lifetime in the absence of either acceptor, and *τ*_*D*(*A*_1_*A*_2_)_ is the donor lifetime in the presence of both acceptors.

The conversion of apparent FRET efficiencies to physical distances is more involved than for two-color FRET due to the additional quenching of the donor by the second acceptor. To define the equivalent of a single-pair FRET efficiency, we ignore the presence of the other acceptor in the rate equation:

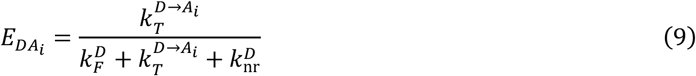

From the definition of the single-pair FRET efficiencies, we calculate the apparent FRET efficiencies for three-color FRET as defined in equation 7:

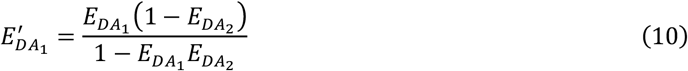

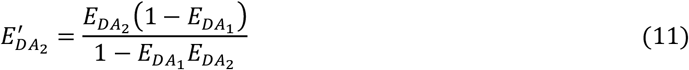

Thus, when the distances are known, the transition probabilities in the three-color FRET system can be calculated by means of equations 1, 10 and 11.

Excitation of the donor dye yields three signal streams and thus two independent intensity ratios, which is insufficient to solve the three-color FRET system for the three inter-dye distances. Thus, to calculate distance-related three-color FRET efficiencies, the FRET efficiency between the two acceptor dyes *E*_*A*_1_*A*_2__ needs to be determined independently. In the experiment, this information is obtained by alternating the excitation of the blue and green dye (Figure 1C-D and Supplementary Figures S1-2). The FRET efficiencies in the three-color FRET system are then calculated using:

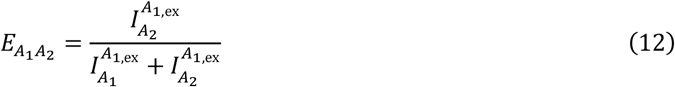

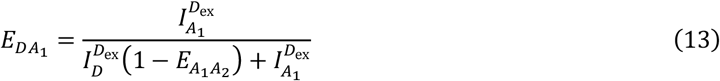

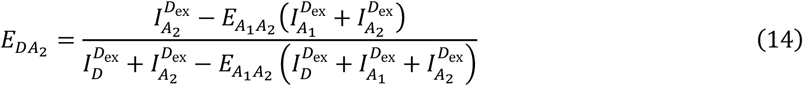

A detailed derivation of equations 10-14 is given in Supplementary Note 1 and may be found in ref. (33). Extensions of equations 13 and 14 with correction factors for real experimental conditions are given in Supplementary Note 2.

In our experiments, a blue, green and red dye are used. Hence, we change the notation from the general case of one donor dye with two acceptor dyes (*D, A*_1_, *A*_2_) to the specific notation indicating the dye colors (*B, G, R*). The emission probabilities *ε* in the three-color FRET system can be calculated according to:

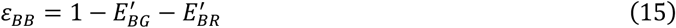

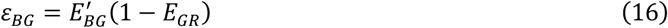

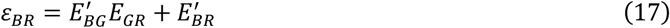

where *ε_ij_* describes the probability to detect a photon in channel *j* after excitation of dye *i*.

These expressions hold true only for an ideal system. As also is the case for two-color FRET, a number of correction factors have to be considered. Additional signal in the FRET channels occurs due to spectral crosstalk of the shorter wavelength dyes into the detection channels of the longer wavelength dyes and direct excitation of dyes by shorter wavelength lasers. Additionally, the different detection efficiencies and quantum yields of the dyes need to be considered. Modified emission probabilities in the non-ideal case are calculated as described in Supplementary Note 3 and 4.

### A likelihood expression for three-color FRET

To extract detailed information about the system in terms of distance heterogeneity, it is necessary to sufficiently sample the FRET efficiency histogram. For three-color FRET, the histogram spans three independent dimensions, requiring the cube of the number of data points as compared to two-color FRET (see Figure S4). In practice, this amount of sampling is usually not accessible. When using the reduced chi-squared 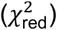 as a determinate for the goodness-of-fit, large counting errors can result in misleadingly small values of 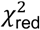 and lead to an over-interpretation of insufficient or bad quality data (see Supplementary Note 5). In such cases, it is advisable to use a maximum likelihood estimator (MLE) to subject the analysis to the likelihood that the observed data was generated by a given model. The MLE approach generally performs better at low statistics and does not require binning of the data (51).

In a three-color FRET experiment, the registered data is a time series of photons in the different detection channels, which are processed into a time-binned set of photon counts {*I*}_*i*_ = {*I_BB_, I_BG_, I_BR_, I_GG_, I_GR_*}_*i*_ with a typical time interval of 1 millisecond. The total likelihood 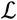 is then the product over the probabilities for all observations that yield the given combination of photon counts {*I*}_*i*_ for a given model *M*:

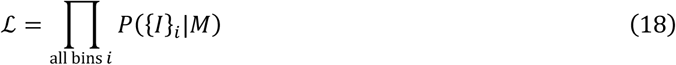

Since excitation of the blue and green dyes is performed alternatingly, both photon emission processes are statistically independent. However, the underlying parameters of the two probability distributions are linked (see equations 15-17). The probability of observing a combination of photon counts after green excitation and after blue excitation is then the product of the individual probabilities:

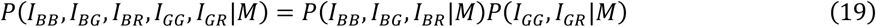

The model *M*, in its simplest form, is described by a triple of distances ***R*** = (*R_BG_, R_BR_, R_GR_*), which are converted into a set of emission probabilities after blue and green excitation ***ε*** = (*ε_BG_, ε_BR_, ε_GR_*) according to equations 1, 16 and 17. To model the FRET processes, a binomial distribution is applied for two-color FRET after green excitation. Analogously, a trinomial distribution is used to describe the three-color FRET process after blue excitation:

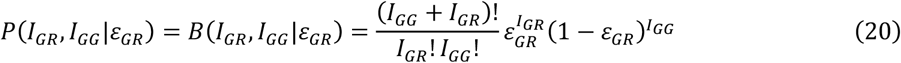

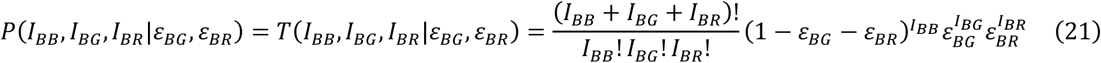

Experimental correction factors are accounted for in the calculation of emission probabilities from distances as described in Supplementary Note 3. Constant background signal in the different detection channels, originating from scattered laser light or detector dark counts, can have a substantial influence on the determined FRET efficiencies, especially in the case of high or low FRET efficiencies where the fluorescence signal in the donor or acceptor channel is low. Since the data is processed into time windows of equal length, background count distributions are assumed to be Poissonian with a mean value of *λ*:

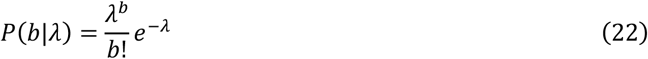

Background contributions are included into the model by summing over all combinations of photon and background counts, whereby the fluorescence signal used to evaluate the FRET processes is reduced by the respective number of background counts.

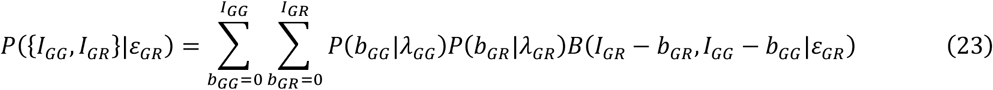

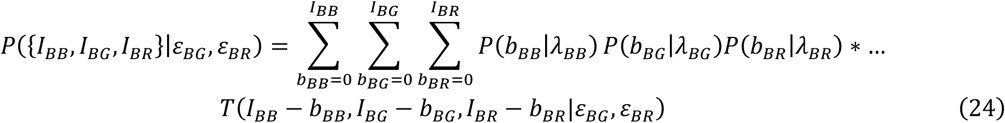

The observed FRET efficiency or proximity ratio histograms usually show broadening beyond the shot noise limit. It has been shown that photophysical artifacts of the acceptor dye can cause additional broadening, which can be described by a Gaussian distribution of distances instead of a single distance with widths of approximately 4-8% of the center distance (18, 52). Broadening beyond this width is typically attributed to conformational heterogeneity of the sample, e.g. due to the existence of conformational substates. To describe additional broadening, a trivariate normal distribution is used to describe a distribution of distances.

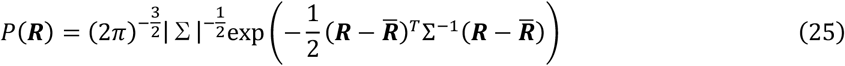

Here, ***R*** = (*R_BG_, R_BR_, R_GR_*) is the column vector of distances, 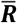 is the column vector of center distances and Σ is the covariance matrix given by:

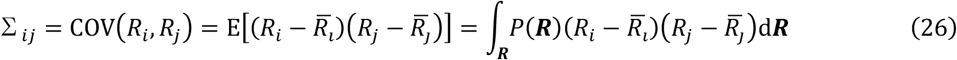

where E[*X*] is the expected value of *X*.

By normalizing the covariance with respect to the individual variances 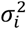, one obtains the Pearson correlation coefficient that quantifies the observed correlation in the interval [−1,1]:

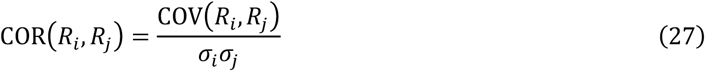

This quantity is a more direct measure of the correlation between the observed distances. The implementation of a Gaussian distribution of distances is achieved by constructing a linearly spaced grid of distances with a given number of bins *G* and width of 2*σ*, thus covering more than 95 % of the probability density, and numerically integrating over the three dimensions:

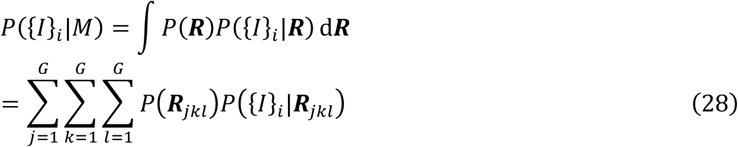

Generally, a grid with *G* = 5 sample points at 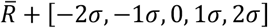 is employed to implement the distance distribution.

Proteins or nucleic acids can exist in multiple conformational states, each of which may show different degrees of static heterogeneity described by a distribution of distances through model *M_j_*. When *N_s_* species are present, the individual likelihoods are summed up and weighted by the normalized contributions *A_j_* of the respective species:

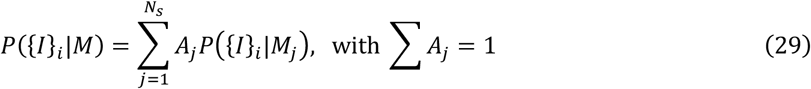

Since the overall likelihood 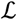 is generally very small, it is necessary to work with the logarithm of the likelihood function to avoid underflow problems:

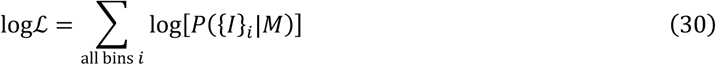

For comparison of the model to the experimental photon count distribution, we use a Monte Carlo approach that simulates the photon emission process to generate expected photon count distributions for the extracted model parameters (for details, see Supplementary Note 6) (19). The Monte Carlo approach is also initially used to find the region of high likelihood (see Supplementary Note 7).

With the likelihood expression for the proposed model of the three-color FRET process, one can find the most likely parameters having generated an observed data set using standard optimization methods. However, since a complex model is applied to a usually limited amount of noisy data, it is of special interest to get a precise idea of how accurately the different parameters are defined given the data at hand. In other words, not only do we want to know the most likely parameters, but also their respective probability distributions, which describe the precision with which we can determine individual parameters. This is a typical problem addressed by Bayesian inference. Using Bayes’ theorem (53), one can relate the probability of certain parameters *θ_i_*, given the observed data {*I*} and background information *α*, to the above derived likelihood by addressing the posterior distribution:

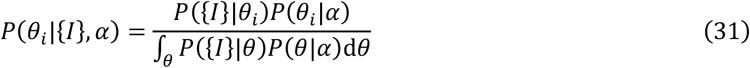

Here, *P*(*θ_i_*|*α*) describes prior information about the parameters derived from the background information *α*. The denominator is a normalization factor, also called the evidence, expressed as the integral of the likelihood over the whole parameter space. Evaluation of the evidence is not feasible in our case as there is no analytical expression available and numerical integration over the whole parameter space is computationally too costly. In Bayesian statistics, however, a number of tools have been developed to estimate the posterior distribution without having to address the evidence. One approach is the Metropolis-Hastings algorithm (47, 48), which samples the posterior distribution by performing random walks over the parameter space using a Markov chain Monte Carlo algorithm. Using independent samples from long random walks (Figure S5A), one can estimate the joint posterior distribution over all parameters and examine marginal posterior distributions to compute the mean and variance of individual parameters (Figure S5B-D). The custom implementation of the Metropolis-Hastings algorithm is outlined in Supplementary Note 8. In the Bayesian inference framework, prior information about the parameters, which may be known from previous experiments, or from the analysis of double-labeled subpopulations, can be incorporated into the analysis. In this way, previous information known about one-dimensional distance distributions can be updated in the Bayesian sense using the triple-labeled molecules to infer additional information regarding the correlation of distances. If no prior information is available, usually a flat prior assigning equal probability over the whole parameter space is used.

## Results and Discussion

### Verification of 3C-PDA using simulations

To test the analysis method, we first applied it to synthetic datasets. Freely diffusing molecules were simulated using a Monte Carlo approach (see Materials and Methods) assuming a molecular brightness of 200 kHz for all fluorophores, similar to what is experimentally observed for our setup. The resulting photon stream was analyzed in the same manner as the experimental data. The simulation assumes fixed distances between the fluorophores resulting in a FRET efficiency of 80% between the green and red dye, 80% between the blue and red dye, and 25% between the blue and red dye. From the photon counts, we calculated three-color FRET efficiencies according to equations 12-14 (Figure 2A-B, grey bars). The conversion of burst-wise photon counts to distance-related three-color FRET efficiencies leads to very broad FRET efficiency distributions with nonsensical values outside the interval [0,1] for the FRET efficiency between blue and red dye. Furthermore, the input value of E_BR_ = 0.25 does not coincide with the peak value in the respective distribution (Figure 2B) and inherent correlations due to stochastic variations in the distribution of photons over the three detection channels are apparent (Figure 2C and S6). Using 3C-PDA, we can determine the underlying inter-dye distances (Table 1) and compare the measured distributions of the FRET efficiencies with expected distributions given by the fitted values (black lines in Figure 2A-C), determined by Monte Carlo simulations (see Supplementary Notes 6 and 7). 3C-PDA accounts for the observed shot-noise broadening of the histograms and accurately captures the apparent correlations in the distributions. To avoid these inherent artifacts, we represent the three-color FRET dataset using the proximity ratios (or signal fractions) PR_BG_, PR_BR_ and PR_GR_ as defined in equations 3 and 5 (Figure 2D-I). To visualize the three-dimensional distribution of occurrences, we show the one- and two-dimensional projections. In this parameter space, the data shows a single population that is well described by the extracted distances. Thus, compared to the transformation into FRET efficiency space, it poses a more natural way of displaying the three-color FRET data. In fact, most single-molecule three-color FRET studies have been analyzed by determining mean values of signal fractions or similar quantities, which are then used to calculate FRET efficiencies and thus distances (35, 36, 40). Using this approach, however, the extracted distances can be very sensitive to errors in the determined mean values (Figure S3), and any information about sample heterogeneity is discarded.

**Figure 2:**
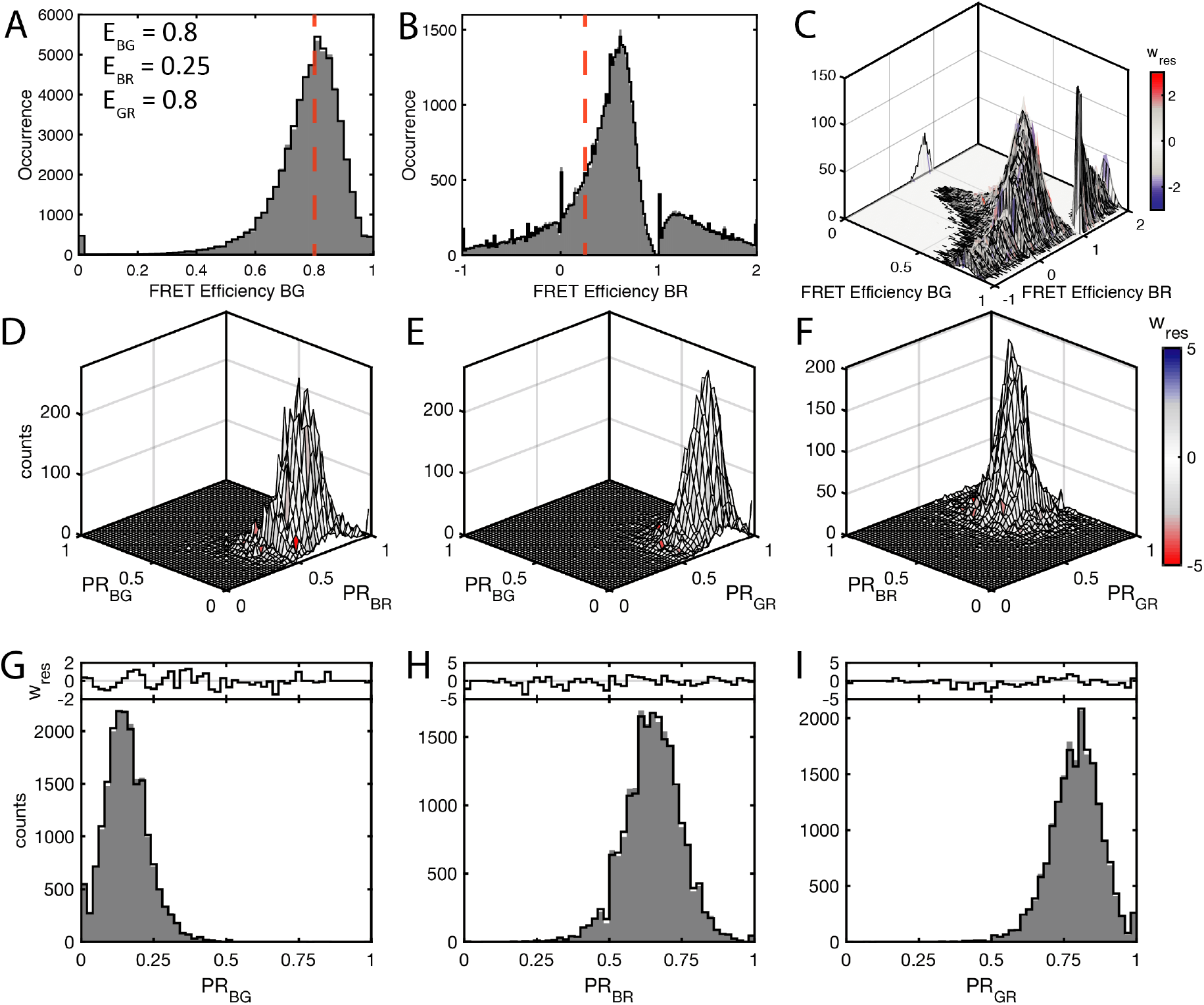
Stochastic noise in three-color FRET. (A-C) Three-color FRET efficiency distributions simulated for an ideal system with the respective two-color FRET efficiencies as listed in (A). The input distances are R_BG_ = 40 Å, R_BR_ = 60 Å and R_BG_ = 40 Å using a Förster distance of R_0_ = 50 Å and a distribution width of *σ* = 2 Å for all dye pairs. FRET efficiencies E_BG_ and E_BR_ show broad and skewed distributions, whereas even values below zero and above one are observed for the FRET efficiency E_BR_. Red dashed lines indicate the expected values of E_BG_ = 0.8 in (A) and E_BR_ = 0.25 in (B). The two-dimensional plot in (C) shows inherent correlations between the two FRET efficiencies due to photon shot noise (compare also Figure S4). Data is shown as grey bars in (A/B) and as a surface plot in (C). Fits are shown as black lines. In (C), the surface used to represent the data is colored according to the weighted residuals. (D-I) Representation of the result of 3C-PDA of the dataset in proximity ratio parameter space. The data is processed into signal fractions after blue excitation (PR_BG_ and PR_BR_) and after green excitation (PR_GR_). Shown are the two-dimensional (D-F) and one-dimensional (G-I) marginal distributions of the three-dimensional frequency distribution. In the two-dimensional projections, the surface representing the data is colored according to the weighted residuals (w_res_). In the one-dimensional projections, the data is shown as grey bars and the 3C-PDA fit as black lines. Weighted residuals are plotted above.

**Table 1:**
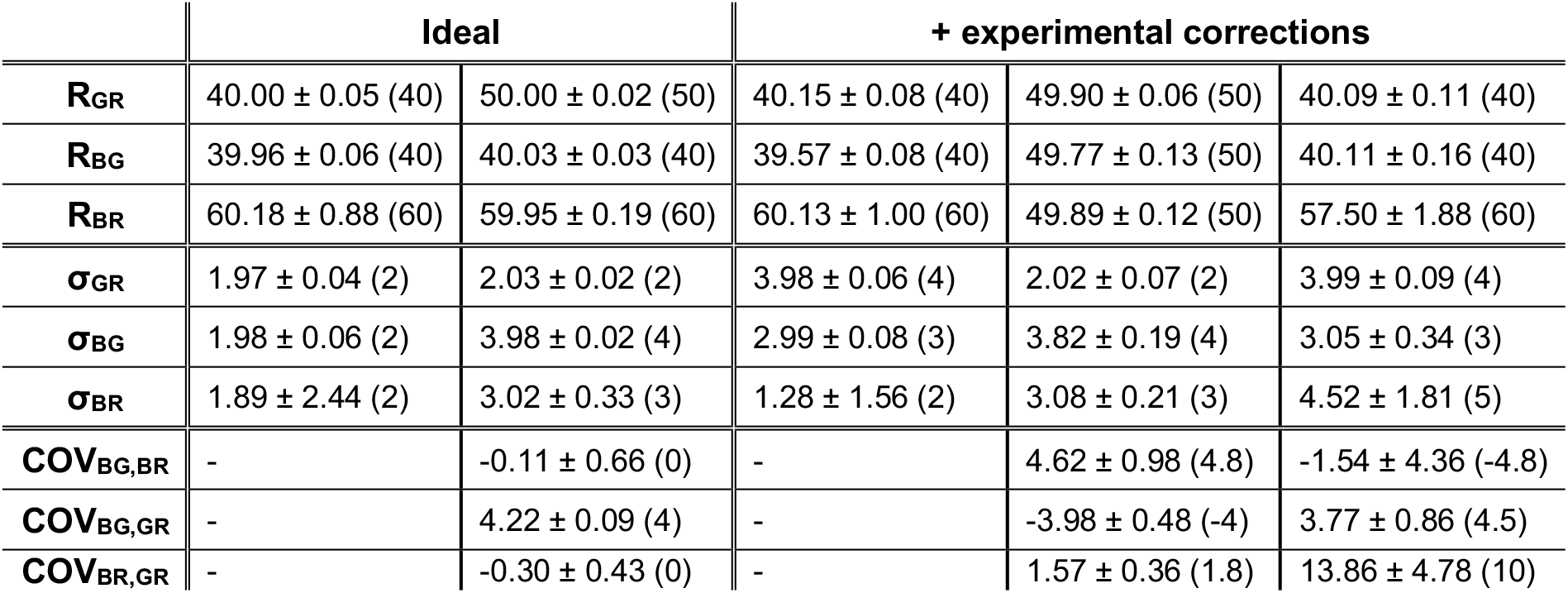
Results of 3C-PDA of simulated data sets with and without experimental correction factors. Distances (R) and distributions widths (σ) are given in Å, while the covariances (COV) are given in Å^2^. Inferred values are shown together with 95 % confidence intervals. Simulation input values are given in brackets. The values used for the experimental correction factors were chosen similar to those obtained experimentally (Table S2). Each analysis was performed on a total number of 10,000 time bins.

To accurately reflect the experimental system, we assumed a normal distribution of distances in the simulations with a width of 2 Å around the center values (Table 1). Using the distance distribution model, we were able to recover the input distances as well as the additional broadening with high accuracy. To map the sensitivity of the method at various inter-dye distance combinations, we simulated different combinations of distances with a constant distance distribution width of 2 Å and a Förster radius of 50 Å for all FRET pairs. To judge the performance at the sampled distances, we determined the precision given by 95% confidence intervals and the accuracy given by the deviation of the inferred values from the input. The precision and accuracy of the analysis depend on the gradient of the signal distribution with respect to the distances. When a change in distance invokes a large change in the distribution of the photon counts over the three detection channels, the inferred values will be well defined. On the other hand, when the photon count distribution is not sensitive to distance changes, the confidence intervals will increase and inaccuracies may occur. To illustrate this effect and to map the sensitive range of 3C-PDA, we plot the fraction of green and red photons after blue excitation as a function of the distances R_BG_ and R_BR_ at a fixed distance R_GR_ of 40 Å (Figure 3A-B). Generally, the sensitivity for the distances R_BG_ and R_BR_ will be optimal when the FRET efficiency between the green and red dye is 0, and worst when the FRET efficiency between green and red is high. In the latter case, the majority of fluorescence signal after excitation of the blue dye is detected in the red channel while only a minor fraction is seen in the green channel, resulting in reduced sensitivity. Mapping the uncertainty at different distance combinations confirms the trend of the signal distributions (Figure 3A-B and Figure S7). The highest uncertainty is observed for the distance R_BR_ in the case of a small distance R_BG_ and a large distance R_BR_, where neither the red nor the green signal fraction shows high sensitivity to a change in the distance R_BR_. However, for the case of R_BG_ = 40 Å and R_BR_ = 60 Å (Table 1), the input values are still recovered with high accuracy and precision.

**Figure 3:**
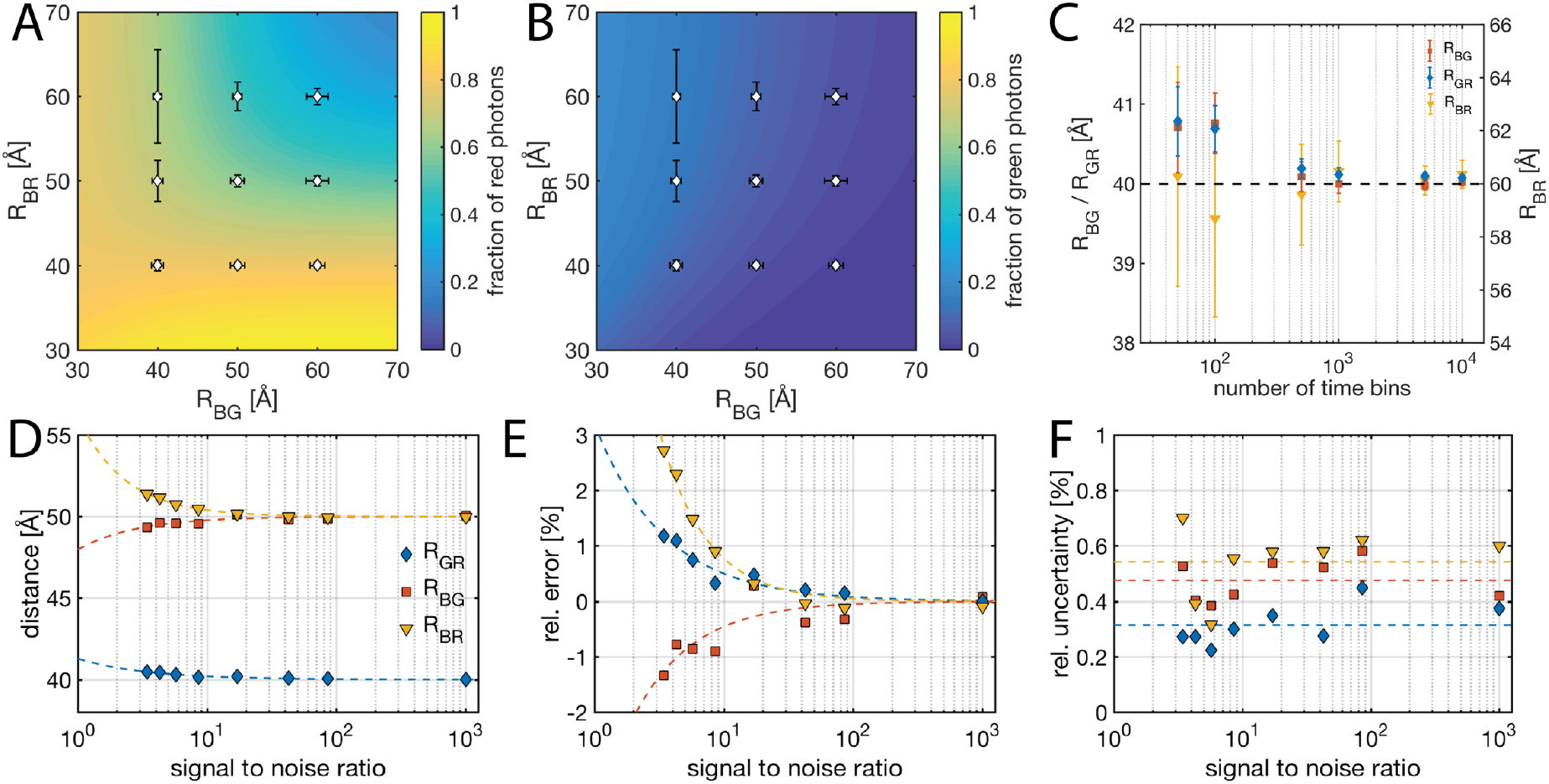
Characterizing the performance of 3C-PDA. (A-B) Map of the distribution of signal after blue excitation on the three-color channels (A: red detection channel, B: green detection channel) at a constant distance between the green and red dye of 40 Å (R_0_ = 50 Å for all dye pairs). Error bars represent the uncertainty at the sampled distance given by the 95% confidence intervals. For better visualization, the absolute uncertainties are multiplied by a factor of 4. Regions where the change in signal is low with respect to distance changes correspond to large uncertainties. See Figure S7 for precision and accuracy maps at different values for R_GR_. (C) Uncertainty of extracted distances given by 95% confidence intervals as a function of the number of time bins used for the analysis. Fitted distance (D), relative error (E) and relative uncertainty (F) as a function of signal-to-noise ratio. To guide the eye, dashed lines, given by power law fits to the data, are shown in D-F.

To address the minimum amount of data needed to accurately infer all three distances, we randomly picked subsets of increasing number of time bins from the simulation using input values of R_GR_ = 40 Å, R_BG_ = 60 Å and R_BR_ = 60 Å, and performed 3C-PDA (Figure 3C). Using 500 time bins, the inferred distances already deviate by less than 2 % from the target values, however the uncertainty is large, especially for the distance R_BR_. At 1000 time bins, the relative error is well below 1 % and the distance uncertainty is reduced to ≤ 2 %, further improving upon inclusion of more time bins. For a simple 3C-PDA including a single population, 1000 time bins are thus sufficient to infer accurate distances. Note that, depending on the duration of the single-molecule observations, one burst can contribute multiple time bins to the 3C-PDA, so the number of sampled molecules can be lower.

To test the sensitivity of our approach with respect to correlated distance broadening, we simulated a static distance distribution with a single non-zero covariance element, using a correlation coefficient of 0.5 between R_BG_ and R_GR_. This corresponds to a system where the blue and red dye are fixed with respect to each other, but the green dye position is flexible, leading to static heterogeneity in the distances with respect to the green dye (Table 1 and Figure S8A-B). When all covariance elements are set to zero, no satisfactory fit could be achieved (Figure S8C). By introducing the covariance matrix as a fit parameter, there is indeed a correlation detected between R_BG_ and R_GR_, while the other distance combinations show no correlation. The analysis yields an inferred correlation coefficient of 0.53 ± 0.01, close to the input value of 0.5 (Table 1 and Figure S8D). Thus, in the ideal case, all input values can be recovered with high confidence, showing that our method is capable of extracting quantitative values for the separation of the three fluorophores as well as the correlation between them.

Experimental systems often coexist in different conformational states. To test to what limit small subpopulations can be detected, we simulated a 9:1 mixture of molecules with a difference in center distance of 2 Å and 5 Å for all distances and a distance distribution width of 2 Å (Figure S9-10). By eye, both datasets can be fit reasonably well using a single population. However, in both cases addition of a second population increases the log-likelihood. To test whether this increase is justified or just caused by the increased number of model parameters, we applied the Bayesian information criterion (BIC) (54) to decide between the two models. The BIC is defined as:

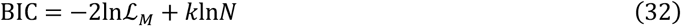

where 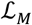 is the maximum value of the likelihood function, *k* is the number of parameters of the model and *N* is the number of data points (here time bins). When comparing different models, the model with the lowest value of the BIC is to be preferred. In this case, the number of model parameters increases from 6 (three center distances and three distribution widths) to 13 (six center distances, six distribution widths and one fractional amplitude). Indeed, the BIC is lower for the two-population model for a separation of 5 Å (Table 2). However, if the distance difference is only 2 Å, the use of the more complex model is not justified by the BIC.

**Table 2:** Testing the detection limit for a minor subpopulation. The main population is defined by center distances of R_GR_ = 60 Å, R_BG_ = 55 Å and R_BR_ = 45 Å with a distribution width of 2 Å. A second population is added at a fraction of 10% with specified distance difference ΔR for all interdye distances. Using the Bayesian information criterion (BIC, see equation 32), the 3C-PDA of the data with one or two populations are compared. For ΔR = 2 Å, the BIC does not justify the use of the model with more parameters (13 versus 6 for a single population). However, a difference of 5 Å justifies the use of the two-population model as indicated by the lower value of the BIC. For a table of all parameters of the analysis, see the Table S1.

| BIC | $\Delta R = 2 \text{ \AA}$ | $\Delta R = 5 \text{ \AA}$ |
| --- | --- | --- |
| 1 population | $1.3072 \cdot 10^5$ | $1.4853 \cdot 10^5$ |
| 2 populations | $1.3075 \cdot 10^5$ | $1.4782 \cdot 10^5$ |
| $\Delta BIC$ | $0.0003 \cdot 10^5$ | $-0.0071 \cdot 10^5$ |

In real experiments, a number of experimental artifacts have to be accounted for. Background noise can lead to an error in the extracted distances, especially if the signal is low. In three-color FRET, this situation can easily occur for the blue channel due to the presence of two FRET acceptors. To test the robustness of our analysis with respect to background noise, we performed simulations using a brightness of 200 kHz for all fluorophores at increasing Poissonian background signal (Figure 3D-F). We define the signal-to-noise ratio (SNR) as the ratio of the average burst-wise count rate over the total background count rate of all detection channels. Typical burst-wise count rates are above 100 kHz and the background count rate per color channel is usually below 2 kHz, resulting in a SNR of 15-20 over all detection channels for our setup. The fitted distances deviate from the input value at low SNR, however at SNR ≥ 10, the relative error is below 1% (Figure 3D-E). The uncertainty of the extracted distances, on the other hand, is independent of the SNR (Figure 3F), since it is primarily limited by the amount of data available.

Additional experimental artifacts are given by crosstalk, direct excitation, and differences in detection efficiency and quantum yield. To test whether we could still infer correct distances in the presence of these artifacts, we included them into the simulations, choosing values close to those encountered in the experiment (Table S1). We were able to still recover the correct center distances for the case of a high FRET efficiency between the blue and green as well as the green and red dyes, and a low FRET efficiency between the blue and red dyes (Table 1). There is, however, a deviation for the inferred value of σ_BR_ and a larger uncertainty associated with both R_BR_ and σ_BR_, likely due to the reduced sensitivity of the red channel caused by the smaller simulated detection efficiency. We then simulated a system showing coordinated broadening of the distance distribution in all dimensions in the presence of experimental artifacts. Mean distances and distribution widths are again accurately recovered, although some deviations can be observed (Table 1). The inferred covariance matrix elements can deviate from simulated values but are within the confidence interval. Revisiting the case of a small distance between blue and green dye and large distance between red and green dye, the existence of substantial coordination can cause deviation of inferred distance values exceeding the inference uncertainty. In this case, the distance R_BR_ is inferred to be 57.5 ± 1.9 Å, while the input value was 60 Å. Likewise, the covariance elements for R_BR_ also deviated from the simulated values, resulting in an underestimation of the anti-correlation between R_BR_ and R_BG_, and an overestimation for the correlation between R_BR_ and R_GR_. Even in this extreme case, the method can thus still recover the correct trend of coordination, albeit the absolute values for the elements of the covariance matrix may deviate from the input values to some extent.

### The likelihood approach to photon distribution analysis

The 3C-PDA method is an extension of previous work on the statistics of photon emission in solution-based two-color FRET experiments (18, 19). By using a likelihood approach, 3C-PDA can be performed without the need of processing the data into histograms. As a result, the Bayesian inference approach is more stable at low statistics, which is of special importance for three-color FRET experiments. We note, however, that our analysis is subjected to pre-processing of the data through burst detection. Thus, determined population sizes will be biased when species differ in brightness because bright bursts are preferentially selected by the burst search algorithm (55). The likelihood analysis presented here may also be applied to two-color FRET experiments. Although acquiring sufficient statistics to sample the proximity ratio histogram usually is not a problem in two-color FRET experiments, we still find that using the likelihood estimator presented here, instead of a *χ*^2^ goodness-of-fit estimator, yields a smoother optimization surface, leading to a faster convergence of the optimization algorithm and generally more accurate results at low statistics. Furthermore, data from two-color FRET measurements of the same distances can be incorporated into the three-color photon distribution analysis to perform a global analysis over many data sets. The two-color FRET data may be taken from incompletely labeled subpopulations, which are available in the three-color FRET measurement, or from independent two-color FRET measurements. In such a global analysis, distances are linked between two- and three-color experiments, however, the dye pairs or setup parameters do not need to be identical for all included datasets. Since the covariance matrix is unique to the three-color FRET dataset, the inclusion of additional two-color FRET datasets increases the robustness of the extracted correlation coefficients between distances. Furthermore, the Bayesian framework presented here allows the natural incorporation of additional information (available e.g. from structural methods such as X-ray crystallography, NMR, cryo-EM or SAXS) into the model by means of the prior probability distribution of the model parameters (see equation 31).

### Application of 3C-PDA to triple labeled DNA

As an experimental benchmark, we measured double stranded DNA labeled with three dyes as a static model system. To see if we could quantitatively detect small changes in a three-color FRET system, we arranged the dyes Alexa488, Alexa568 and Alexa647 such that significant FRET could occur between all dye pairs. The green and red dyes were positioned on one strand of the double helix at a distance of 27 bp, while the blue dye was positioned on the complementary strand between the two acceptor dyes. Two constructs were designed where the position of the blue dye was shifted by 3 bp, resulting in distances of 17 bp to the green dye and 10 bp to the red dye for construct 1 (DNA-Alexa1), and 14 bp to the green dye and 13 bp to the red dye for construct 2 (DNA-Alexa2) (Figure 4G). The 3C-PDA fit of DNA-Alexa1 is shown in Figure 4A-F. While not directly apparent from the photon count distributions, a second population with ~20% contribution was required to describe the data. DNA-Alexa2 more visibly showed a second population with a lower value for PR_BG_, accounting for ~15% of the measured data (Figure S11B). Here, we focus the analysis on the main population (Table 3 and Figure 4H) and attribute the minor secondary population to a photophysical artifact of the dyes, discussed further below. As expected, the distance between the green and red dye is unchanged for both arrangements at 94.9 Å and 96.4 Å for DNA-Alexa1 and DNA-Alexa2, respectively. Changing the position of the blue dye results in a distance change for R_BG_ from 64.4 Å to 52.3 Å, and for R_BR_ from 62.2 Å to 72.6 Å, resulting in a difference of ΔR_BG_= −12.1 Å and ΔR_BR_= 10.4 Å. To investigate whether we could infer correct distances, we compare our result with expected distances as determined by accessible volume (AV) simulations (56). When comparing the relative distance differences between the two constructs, we find very good agreement between measurement and AV simulations, which yield ΔR_BG_= −13.5 Å and ΔR_BR_= 10.8 Å (Table 3). The absolute inter-dye distances for R_GR_ and R_BG_ also match reasonably well with the AV-derived distances, while a large deviation is observed for R_BR_ (*R*_exp_–*R*_AV_ = 13/18 Å for DNA-Alexa1/2, Table S3). We separately analyzed double-labeled subpopulations of the same measurement using 2C-PDA (Table S3 and Figure S12) and performed measurements on DNA molecules carrying two of the three dyes (Table S3 and Figures S13 A-D). The results agree well with the distances determined by 3C-PDA, showing that the deviation of absolute distances from expected values for R_BR_ is not an artifact of the 3C-PDA method (see Supplementary Notes 9 and 10 for further discussion of potential photophysical and structural artifacts).

**Figure 4:**
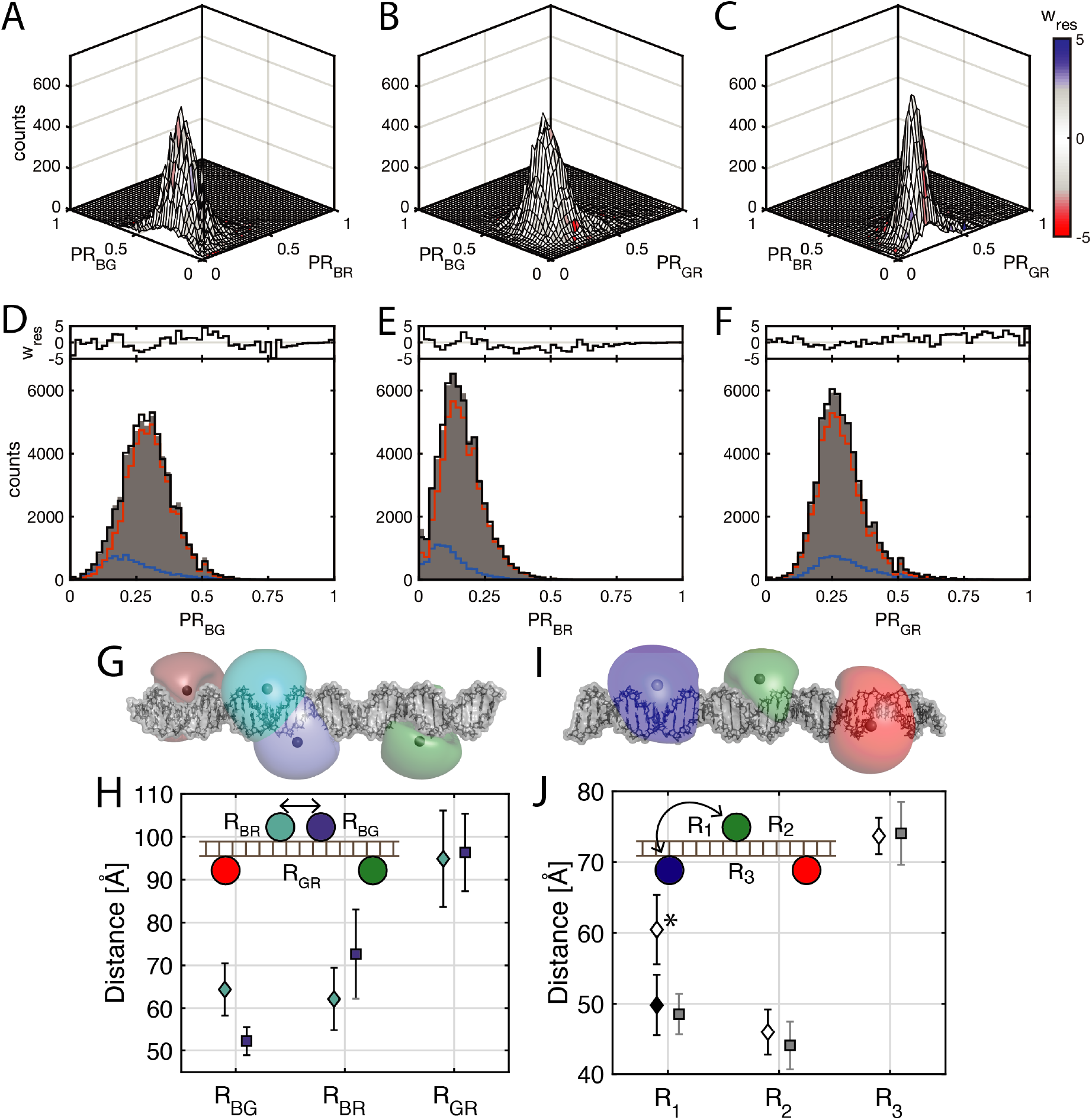
3C-PDA of triple-labeled DNA. (A-F) Exemplary fit result of 3C-PDA of DNA-Alexa1, corresponding to the cyan position of the blue dye in the structural model of the dsDNA construct shown in panel G. Shown are the projections of the three-dimensional histogram of signal fractions PR_GR_, PR_BG_ and PR_BR_. Two-dimensional projections are displayed as surface plots in the top row (A-C). The surface is colored according to the weighted residuals as indicated by the color bar. In the bottom row (D-F), one-dimensional projections (grey bars) are plotted together with the fit result (black line). The two components of the model function are shown as red and blue lines. Weighted residuals are shown above. All DNA constructs required the use of two Gaussian distance distributions to describe the data. The 3C-PDA of DNA-Alexa2 and DNA-Atto1/2 are shown in Figure S11 and S15, respectively. (G) Structural model of the DNA-Alexa constructs. The accessible volumes of the dyes Alexa488 (cyan: DNA-Alexa1, blue: DNA-Alexa2), Alexa568 (green) and Alexa647 (red) are shown as clouds. Spheres with the respective colors indicate the mean positions of the dyes. (H) Fit results for DNA-Alexa measurements. Cyan diamonds show the results with the blue dye in the left position (DNA-Alexa1, cyan cloud in the structure above), whereas blue squares indicate the blue dye in the right position (DNA-Alexa2, blue cloud). Fitted distribution widths are given as error bars. The change of the position of the blue dye by 3 bp results in an anticorrelated change of the distances RBG and RBR. (I) Structural model of the construct DNA-Atto1. The accessible volumes of the dyes Atto488 (blue), Atto565 (green) and Atto647N (red) are shown as clouds. Gray spheres indicate the mean positions of the dyes. In the DNA-Atto2 construct, the positions of the blue and green dyes are switched. (J) Fit results for the DNA-Atto constructs. Diamonds indicate the results of DNA-Atto1, corresponding to the arrangement of dyes as shown in the inset. Gray squares indicated the arrangement where the blue and green dye positions have been exchanged (DNA-Atto2). Switching of the positions of blue and green dye should not affect the recovered distances. This holds true for distances R2 and R3. For the distance R1, however, a large difference is observed (*). This deviation originates from dynamic quenching of Atto565 in construct DNA-Atto1 (see main text and Supplementary Note 9). By accounting for the quenching in the analysis, the correct distance can still be recovered for R1 in DNA-Atto1 (black diamond). Error bars represent the fitted distribution widths.

**Table 3:**
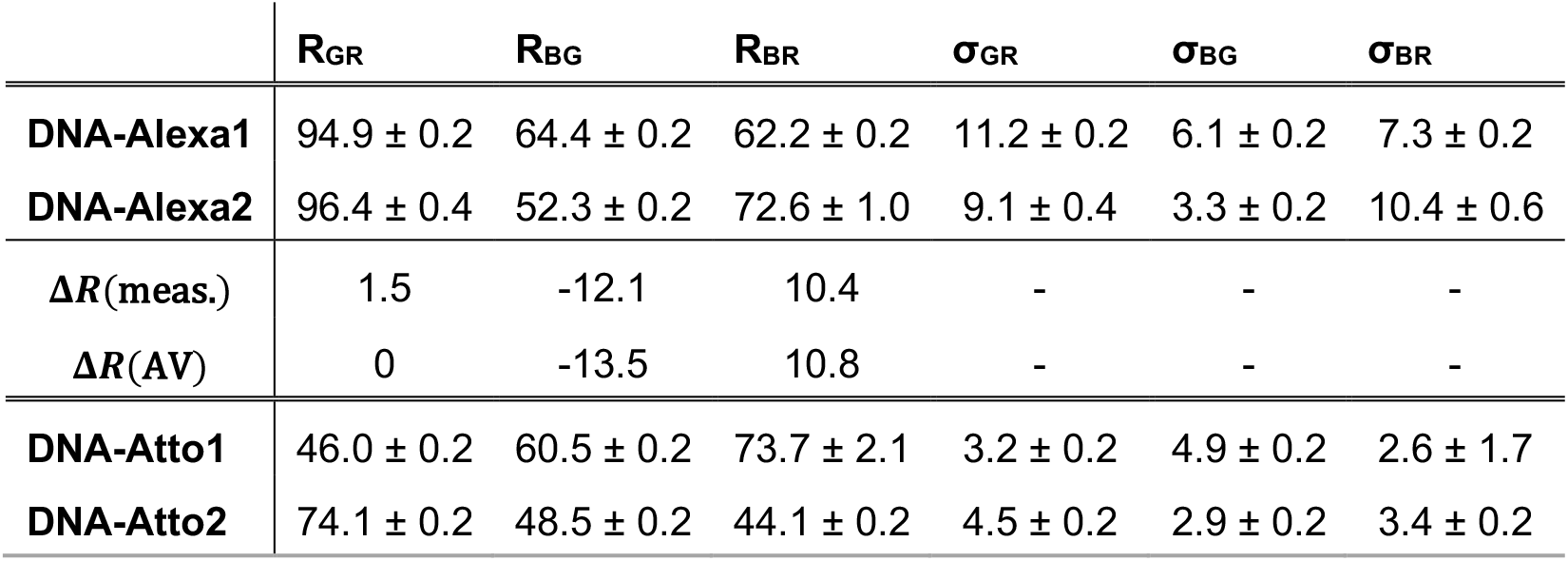
Results of the 3C-PDA of DNA constructs. Distances determined by 3C-PDA of the DNA constructs labeled with Alexa488, Alexa568 and Alexa647 (DNA-Alexa1/2) or with Atto488, Atto565 and Atto647N (DNA-Atto1/2). The inferred distances for the major population (>80% contribution) are listed with associated inference uncertainties given as 95% confidence intervals. For DNA-Alexa measurements, experimental distance changes, *ΔR*(meas.), are compared to theoretical distance changes as determined from accessible volume (AV) calculations, *ΔR*(AV). For a complete list of all inferred model parameters, comparison to two-color control constructs and theoretical distances estimated from accessible volume calculations, see Table S3-4.

We then tested another combination of fluorophores by arranging the dyes Atto488, Atto565 and Atto647N first in a cascade geometry (Figure 4I). This arrangement (DNA-Atto1) is expected to show a high FRET efficiency from the blue to green dye and the green to red dye, while the FRET efficiency from the blue to red dye will be very low, similar to the situation studied using simulated datasets in the previous section. This resulted in similar distributions of the three-color FRET efficiencies as observed for the simulated data, including the occurrence of false correlations (Figure S14). A single population was again not sufficient to describe the data. Therefore, a second population was included in our fit, accounting for approximately 15% of the data (Figure S15A and Table S4). For the main population, we find a large distance of 74 Å between the blue and red dye and shorter distances between the green and red dye (46 Å) as well as the blue and green dye (61 Å) (Table 3 and Figure 4J). We then switched the positions of the blue and green dye such that substantial energy transfer could occur between all dyes (DNA-Atto2). Again, inclusion of a small secondary population was necessary, accounting for 20% of the data (Table S4 and Figure S15B). As expected, the distance between the green and red dye is now similar to the distance between the blue and red dye in the cascade arrangement (74 Å). Likewise, the distance between the blue and red dye in DNA-Atto2 is close to the distance measured between the green and red dye in DNA-Atto1 (44 Å). Surprisingly, we find a significant disagreement for the distance between the blue and green dye at 48 Å, which should be identical in both arrangements (indicated by a * in Figure 4J). We analyzed separate measurements of the same constructs carrying only two of the three dyes by 2C-PDA, which resulted in similar distances for DNA-Atto2 as compared to the 3C-PDA results (Figure S13 E-H and Table S4). In DNA-Atto1, however, the dye Atto565 showed position-dependent quenching causing the observed deviation for the distance R_BG_. The quenching likely occurs due to photoinduced electron transfer to nearby guanine residues, as has previously been reported for rhodamine-based fluorophores (61) (see Supplementary Note 11 for a detailed discussion). By selective analysis of the unquenched population using 2C-PDA, we can recover comparable values for R_BG_ for both constructs, yielding a corrected value for DNA-Atto1 of ~50 Å (black diamond in Figure 4J). Such a selective analysis was not possible for the three-color FRET data because the fluorescence lifetime of Atto565 was already significantly reduced due to the high FRET efficiency to the red dye (E_GR_ > 0.9, *τ_G_* < 1 ns), making it difficult to separate quenched and unquenched populations.

In our model function, we assumed a Gaussian distribution of inter-dye distances. A Gaussian distance distribution model has been standard for almost all PDA studies so far, even for systems that are expected to be rigid such as dsDNA (18, 19). A detailed discussion of the origin of the broadening of the FRET efficiency histograms beyond the shot-noise in the absence of conformational heterogeneity is given by in reference (52) and is attributed to the existence of different photophysical states of the acceptor leading to an apparent distribution width that is proportional to the mean interdye distance. The observed distribution widths in the 3C-PDA of dsDNA are in the range of 8.4 ± 3.1% of the center distances (averaged over all dye combination). This value is comparable to previously reported results for two-color PDA analyses of dsDNA labeled with Alexa488 and Cy5, yielding a value of 7.6% (52). Here, we did not consider the covariances in the analysis of the DNA constructs. When artifacts due to dye photophysics are the main contribution to the width of the observed distance distribution, e.g. the existence of multiple photophysical states, 3C-PDA will reveal apparent false correlations. Considering, for example, a quenched state of the red dye, the distances to both the green and the blue dye would be affected, which would result in a false-positive covariance for these distances. While it is in principle possible to use the off-diagonal elements of the covariance matrix to investigate the correlation between distances, special care has to be taken when investigating the covariance matrix in terms of physically relevant conformational broadening. Moreover, whereas artifacts originating from the presence of multiple photophysical states can usually still be described by a single population with an increased distribution width in 2C-PDA, 3C-PDA is more sensitive to these artifacts. In this case, the covariance matrix of the distance distribution would be required to describe the correlated broadening of the apparent distribution of distances. As such, it is not unexpected that two populations were required to describe the three-color FRET data in all cases. Indeed, the minor secondary population observed in the DNA measurements most likely originates from photophysical artifacts such as spectral shifts of the fluorophores reported for Atto647N (62, 63) and Alexa488 (64, 65) or sticking interactions of the dyes to the DNA surface (52, 60).

### Towards quantitative three-color FRET in proteins

To test 3C-PDA on a protein system, we labeled the Hsp70 chaperone BiP (binding immunoglobulin protein) with the dyes Atto488, Atto565 and Atto647N (Figure 5A). During its nucleotide-dependent conformational cycle, BiP undergoes a large conformational change whereby the lid of the substrate binding domain assumes an open conformation when ATP is bound but is closed in the ADP-bound state (23, 29). By positioning the green and red dyes on the flexible lid and the substrate-binding domain (SBD), we can monitor the state of the lid (open/closed). The blue dye is specifically attached to the nucleotide-binding domain (NBD) through introduction of an unnatural amino acid. The labeling of the green and red dye, however, was performed stochastically using two site-specifically introduced cysteines, resulting in a random distribution of the green and red dye labels between the two possible configurations BGR and BRG. Under the assumption that the environment of the dyes is similar at both labeling positions, the measured distance between the green and red dyes should not be affected. In 3C-FRET, however, a superposition of the two configurations is observed. The model function can be extended to account for the stochastic labeling using only one additional parameter, which is the relative population of the two configurations BGR and BRG. Considering that a permutation of the positions of green and red dye is equivalent to adding a second population with interchanged distances BG and BR, the total likelihood per time bin is then given by:

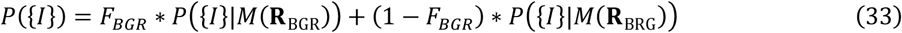

where F_BGR_ is the fraction of molecules with labeling scheme BGR. When the labeling positions are not equally accessible for the two acceptor dyes, the resulting distribution will be uneven. In such cases, where F_BGR_ significantly differs from 0.5, it can be extracted from the fit as a free fit parameter. A more robust analysis can be performed when F_BGR_ is determined by other methods (e.g. by mass spectrometry).

**Figure 5:**
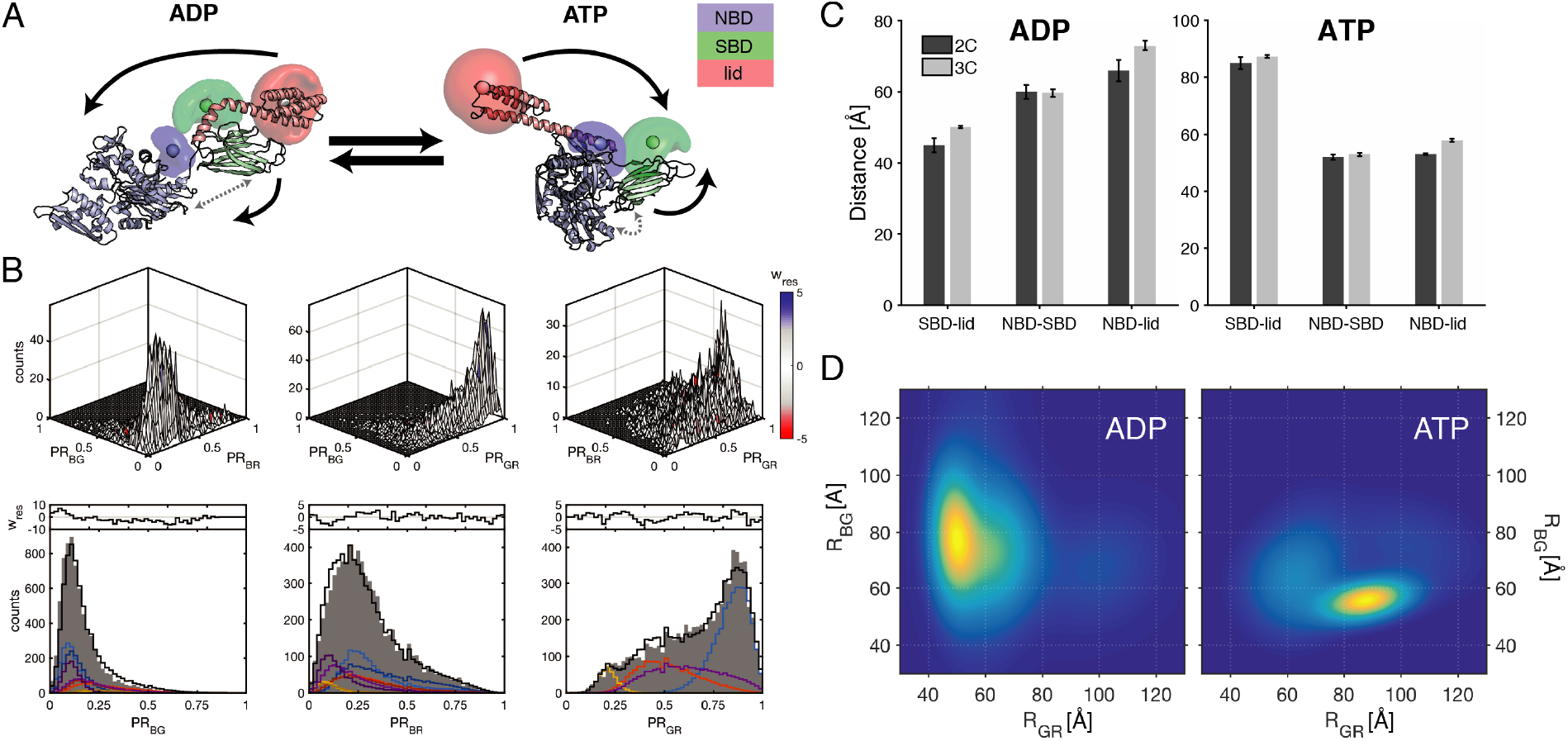
3C-PDA of the heat shock protein BiP. (A) Structural representations of the Hsp70 chaperone BiP in the ADP-bound state (left, based on the crystal structure of the homolog DnaK, PDB: 2KHO) and in the ATP-bound state (right, PDB: 5E84). The nucleotide-binding, substrate-binding and lid-domain (NBD, SBD and lid) are color-coded in blue, green and red respectively. Dye positions are indicated by clouds that represent the possible positions determined using accessible volume calculations. The lid of the SBD is closed in the ADP-bound state but opens up in the ATP-bound state. Black arrows indicate the movement of domains during the nucleotide-dependent conformational cycle. The distance between the SBD and NBD is larger in the ADP-bound state, while the two domains directly contact each other in the ATP-bound state (grey arrows). (B) 3C-PDA of BiP in the ADP state. Shown are the projections of the three-dimensional histogram of signal fractions PR_GR_, PR_BG_ and PR_BR_. Two-dimensional projections are displayed as surface plots in the top row. The surface is colored according to the weighted residuals as indicated by the color bar. In the bottom row, one-dimensional projections (grey bars) are plotted together with the fit result. Weighted residuals are shown above the plots. The individual components of the fit are shown in blue, red, yellow and purple in the onedimensional projections. Additionally, the second population originating from stochastic labeling is shown in darker color (dark blue, dark red, dark yellow, dark purple) in the one-dimensional projections of PR_BG_ and PR_BR_. (C) Comparison of distances extracted from two-color FRET experiments of BiP (dark grey), taken from Rosam et al. (29), and determined using 3C-PDA (light grey). D) Two-dimensional distance distributions extracted from the measurements of BiP in the presence of ADP (left) and ATP (right) using 3C-PDA without correcting for the stochastic labeling. The distance between the green and red fluorophores, R_GR_, reports on the distance between the lid-domain and the substrate-binding domain. The distance between the blue and green fluorophores, R_BG_, monitors the distance between the substrate-binding and the nucleotide-binding domain. The 3C-PDA confirms the picture obtained from the crystal structures (A): In the presence of ADP, the lid is closed and the interdomain distance shows a broad distribution, indicative of conformational heterogeneity; in the presence of ATP, the lid closes and the distribution width of the interdomain distance is narrower, showing a more defined conformation. See Fig. S18 for the associated 3C-PDA fits and Fig. S19 for a complete display of the inferred distance distribution.

Here, we performed 3C-PDA on BiP in the presence of ADP and ATP. The conformational cycle of BiP has previously been studied by single-pair FRET using the identical labeling positions (23, 29). We determined the degree of labeling for the individual dyes using absorption spectroscopy, yielding an upper estimate for the triple-labeling efficiency of ~25 %. However, after filtering for photobleaching and -blinking, the fraction of usable single-molecule events for 3C-PDA was reduced to less than 5% of all detected molecules. For the ADP-state, the obtained proximity ratio histograms show a broad distribution with a peak at high proximity ratio GR and low proximity ratio BG and BR (Figure 5B). The ATP-state, on the other hand, is characterized by a main population at low proximity ratio GR and BR, whereas the proximity ratio BG shows a bimodal distribution (Figure S16). Inspection of the one-dimensional projections does, however, not reveal the full complexity of the data, which is more evident in the two-dimensional projections. As such, it is not straightforward to assign the number of populations based on the one-dimensional projections alone. To fit the data, we assumed the occupancy of the two cysteine labeling positions to be equal between the green and red dye (*F_BGR_* = 0.5). At a minimum, we required three components to describe the data. However, since inclusion of a fourth component increased the likelihood and decreased the BIC (Table S5), we fit the data using a total of 4 populations (Figure 5B, S16 and Table S6). Without further knowledge about the distribution of the green and red labels, the three-color FRET distances R_BG_ and R_BR_ cannot be assigned to physical distances in the molecule. However, by reducing the analysis to the main population (colored blue in the figures), we can assign the three-color FRET derived distances by simple comparison with the previously determined values using single-pair FRET (Figure 5C) (29). Reasonable agreement is obtained between two-color and three-color FRET experiments for all distances, showing that we can follow the conformational cycle of BiP with three-color FRET.

Previously, the observed two-color FRET efficiency distributions for the NBD-lid and NBD-SBD FRET sensors (i.e. the distances between the blue/red and the blue/green labels in Figure 5A) were found to be very similar (23, 29). Based on this observation, we also performed a simplified analysis using a four-component model without accounting for the stochasticity of the fluorescent labeling (Figure S17). Indeed, also the inferred distributions from 3C-PDA for R_BG_ and R_BR_ are very similar (Figure S18 and Table S7), indicating that both distances may be used interchangeably to address the interdomain distance. The inferred two-dimensional distribution of the SBD-lid (R_GR_) and SBD-NBD (here R_BG_) distances are shown in Figure 5D (see Figure S18 for all distance pairs). From the main populations of the distance distribution, we can confirm the picture obtained from the crystal structures (Fig. 5A), showing that the protein indeed undergoes a coordinated conformational change. In the presence of ADP, the lid-domain is closed (R_GR_ ≈ 50 Å, σ_GR_ ≈ 5 Å), while the interdomain distance shows a broad distribution indicating conformational heterogeneity (R_BG_ ≈ 80 Å, σ_BG_ ≈ 15 Å, Fig. 5D, left). In the ATP-state the lid opens up (R_GR_ ≈ 89 Å, σ_GR_ » 10 Å), while the domains come into closed contact, resulting in a narrower distribution width for the interdomain distance (R_BG_ ≈ 56 Å, σ_BG_ ≈ 5 Å, Fig. 5D, right). The simplified model thus allowed us to characterize the conformational space of BiP using three-color FRET and investigate correlated distance changes from the inferred multidimensional distance distributions.

While we generally expected broad distributions for the three-color FRET data, we also observed large conformational heterogeneity for the sensor between the substrate-binding domain and the lid, which previously was found to adopt more defined conformations (23, 29). A possible explanation for the excess heterogeneity might be given by the contribution of contaminations. Through the filtering for triple-labeled molecules that showed no photo-bleaching, we are subjecting our analysis to a small fraction of generally less than 5% of all detected molecules. Thus, it is expected that fluorescent impurities, which by chance pass the filtering process, may have a higher contribution compared to two-color FRET experiments. Although BiP was still functional after the triple-labeling step with respect to its ATPase activity (see Figure S19), we also cannot exclude that the labeling with three dyes might slightly destabilize the structure of the protein.

Biomolecules generally often show complex conformational landscapes. Here, we required a total of four populations to describe our data. Due to the sensitivity of 3C-PDA to correlations in the data, it is expected that more states are necessary to fully describe the data as compared to two-color FRET experiments. On the other hand, the sensitivity of 3C-PDA is also what makes it possible to reliably extract more information from the data. Using the example of the Hsp70 chaperones, previous studies suggested that the opening and closing of the lid correlates with the distance between substrate-binding and nucleotide-binding domains (66, 67). However, each state of the lid (open/closed), might show multiple states of the interdomain sensor. Thus, while two-color experiments on the SBD-lid sensor could be well described by two states, and the interdomain sensor fit well to a broad distance distribution, three-color experiments naturally require a larger number of states due to the multidimensional and correlated information available. With respect to the Hsp70 chaperones, the question of the allosteric communication between the nucleotide-binding and substrate-binding domains is a key question to understand their function (66, 68). 3C-PDA is a promising method to elucidate the coordination between substrate binding and nucleotide hydrolysis within these proteins. Fundamentally, our results on the conformational changes of BiP are limited by the lack of specific labeling for the two acceptor dyes in the experiment. We describe how this limitation can be overcome when the labeling efficiencies of the acceptor positions are different and known *a priori*. Promisingly, many novel approaches for the specific labeling of proteins have been developed in the past years (44–46), enabling the attachment of three or more fluorophores to single proteins through a combination of orthogonal labeling approaches. Moreover, it is worth noting that 3C-PDA is, of course, not limited to the study of intramolecular distances in single proteins. Rather, it has promising applications in the study of interactions between proteins and/or nuclei acids, whereby the interaction partners may be labeled with a single or two fluorophores each.

## Summary and Conclusions

Here, we presented 3C-PDA, a statistical method for the analysis of single-molecule three-color FRET burst analysis experiments. The method incorporates the underlying physical distance heterogeneity into the model function, enabling a detailed description of the conformational space of biomolecules by three distances simultaneously. The need to apply statistical methods to the analysis of three-color FRET data arises due to the inherent noise that occurs when converting the limited number of photons per molecule to three-color FRET efficiencies. Our model-based Bayesian inference approach describes the observed data with a distance distribution model that includes all needed experimental correction factors, thus presenting an effective way to de-noise three-color FRET experiments and extract information about the underlying distance heterogeneity. The experimental results can be directly interpreted in terms of distance changes, using the three-color FRET information to reveal the existence of coordinated conformational changes. The theory described here applies to systems that show no conformational dynamics on the millisecond time scale. The framework can, however, be extended to account for dynamic interconversion between distinct conformational states, which will enable the study of complex dynamic processes within single biomolecules.

## Materials and Methods

### Simulations

Simulations of single-molecule three-color FRET experiments were performed using a Monte Carlo approach. Diffusion of molecules in a box of size 7 μm × 7 μm × 12 μm was simulated by drawing normally distributed random numbers for the displacement at each time step Δ*t* of 1 μs with a width given by 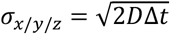. Molecules exiting the box were re-inserted on the opposite side (periodic boundary condition). The observation volume was represented by a 3D Gaussian function with lateral and axial width of 500 nm and 1500 nm at 1/*e*^2^ of the maximum intensity, respectively. Photons for each excitation channel were generated randomly at each time step with a probability proportional to the excitation intensity at the current position. FRET and experimental artifacts were evaluated on photon emission as described in Supplementary Note 3. Simulation parameters were chosen to approximately represent experimental conditions with a diffusion coefficient of 125 μm^2^/s and a brightness of 200 kHz at the center of the confocal volume. To simulate static broadening of distance distributions, a new set of distances was drawn according to the distance distribution for each molecule every 10 ms.

### Experimental Setup

Three-color FRET experiments with pulsed interleaved excitation (PIE) (15, 69) and multiparameter fluorescence detection (MFD) (8) were performed on a homebuilt confocal three-color dual-polarization detection setup (Figure S1) based on a Nikon Eclipse Ti-DH inverted microscope. For pulsed interleaved excitation, the three lasers (LDH-D-C-485, LDH-D-TA-560, LDH-D-C-640, PicoQuant, Berlin, Germany) are synchronized by a laser driver (Sepia II, PicoQuant) at a frequency of 16.67 MHz with 20 ns delay between consecutive pulses to minimize temporal crosstalk between PIE channels. The lasers are combined into a polarization maintaining single-mode fiber (QPMJ-A3A 405/640, OZ Optics, Ottawa, Canada), collimated (60SMS-1-4-RGBV11-47, Schäfter+Kirchhoff, Hamburg, Germany) and focused into the sample by a 60x 1.27 NA water immersion objective (Plan Apo IR 60x 1.27 WI, Nikon, Düsseldorf, Germany). The average excitation power measured before the objective was 100 μW, 90 μW and 70 μW for the blue, yellow and red lasers, respectively. Fluorescence was separated from the excitation light by a polychroic mirror (zt405/488/561/633, AHF Analysentechnik, Tübingen, Germany) and focused through a 50 μm pinhole. The signal was then split into parallel and perpendicular polarization with respect to the excitation by a polarizing beam splitter (Thorlabs, Dachau, Germany) and spectrally separated into the three spectral channels by two dichroic mirrors (BS560 imaging, 640DCXR, AHF Analysentechnik) and three emission filters per polarization (ET525/50, ET607/36, ET670/30, AHF Analysentechnik). Photons were detected using six single-photon-counting avalanche photodiodes (2x COUNT-100B, LaserComponents, Olching, Germany, and 4x SPCM-AQR14, Perkin Elmer, Waltham, Massachusetts) and registered by TCSPC electronics (HydraHarp400, PicoQuant), which was synchronized with the laser driver.

### DNA sample preparation and measurement

Single- and double-labled ssDNA was purchased from IBA GmbH (Göttingen, Germany). DNA was labeled either with Atto488, Atto565 and Atto647N (Atto-Tec, Siegen, Germany) or Alexa488, Alexa568 and Alexa647 (Thermofisher Scientific). DNA sequences and labeling positions are given in the SI. Single-stranded DNA was annealed in TE buffer (10 mM Tris, 1 mM EDTA, 50 mM NaCl, pH 8.0) by heating to 95 °C for 5 min and then gradually cooling down at a rate of 1 °C per minute to a final temperature of 4 °C. The sample was diluted to a final concentration of 20 pM in PBS buffer (Thermofisher Scientific) containing 1 mM Trolox (Sigma Aldrich) to reduce photobleaching and -blinking of the dyes(70).

### Data analysis

Data processing was performed using the software package *PAM* written in MATLAB (The MathWorks, Natick, Massachusetts), which is freely available (see Code Availability) (71). Bursts were selected as described previously using a sliding time window approach on the total signal, requiring at least 5 photons per time window of 500 μs and at least 100 photons in total per burst (19). Triple-labeled bursts were selected based on stoichiometry thresholds (S_BG_, S_BR_ and S_GR_) (35) using a lower boundary of 0.15 and an upper boundary of 0.9. To additionally remove photobleaching and blinking events, the ALEX-2CDE filter (72) was applied, which was calculated pairwise for the three excitation channels. Fluorescence lifetimes and anisotropies for the PIE channels BB, GG and RR were determined as described previously (17). For the photon distribution analysis, burstwise data was processed into time bins of 1 ms length. Using three-color MFD-PIE, we can determine all necessary correction factors directly from the experiment, with the exception of the direct excitation probability for the PDA, which is different from the direct excitation correction factor used to correct photon counts. Direct excitation correction of FRET photon counts was performed based on the signal from the alternating acceptor excitation (see Supplementary Note 2), whereas, for the PDA, one requires the relative probability that the donor excitation laser excites the acceptor dye (see Supplementary Note 4). Crosstalk is determined from single-dye species, which are isolated using the sorting capabilities available in three-color PIE-MFD. *γ*-factors, accounting for the differences in quantum yield and detection efficiency for the different dyes, can be determined using the lifetime information by plotting the FRET efficiency as determined from photon counts versus the donor lifetime and minimizing the deviation from the theoretical static FRET line (20). This method is applicable only when the measured system shows no dynamics. For two-color FRET, the static FRET line is given by:

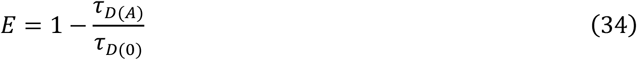

where *τ*_*D*(0)_ and *τ*_*D*(*A*)_ are the donor dye lifetimes in the absence and in the presence of the acceptor dye. An analogous relation is found for the total FRET efficiency from the blue dye to both acceptor dyes as shown in equation 8. In the case of flexible linkers, the static FRET line has to be modified to account for the distance averaging occurring over the burst duration (20). To apply this method to three-color FRET, we extended the theory to determine the static FRET line in the case of linker fluctuations as described by Kalinin et al. (20) for a three-color FRET system (see Supplementary Note 12). Since the systems discussed here are static, *γ*-factors can be extracted directly from the experiment. For three-color FRET, three *γ*-factors have to be considered, defined as the ratio of quantum yield *Φ* and detection efficiency *g* of the two dyes by 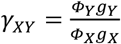. The three *γ* -factors are approximately related by *γ_BR_* = *γ_BC_* ∗ *γ_GR_*. To determine the *γ*-factors for a three-color FRET measurement, first *γ_GR_* is determined by minimizing the deviation from the static FRET line as described above. Subsequently, the deviation of *E*_*B*→*G*+*R*_ from the static FRET line with respect to the lifetime of the blue dye is minimized by varying either *γ_BG_* or *γ_BR_*. The third *γ*-factor is then calculated using the above-mentioned relation between the *γ*-factors. We verified this approach by separate measurements on double-labeled DNA constructs with two FRET species, where we determined the *γ*-factor using the relation between FRET efficiency and stoichiometry, which, within error, yielded equivalent values (Table S2).

### Determination of confidence intervals

Inference uncertainties are given by 95% confidence intervals determined from long runs of the Metropolis-Hastings algorithm to estimate the posterior probability density (> 10.000 steps). Statistically independent samples are drawn with a spacing of 500 steps and parameter uncertainties are determined from the distribution of samples.

### Protein constructs

For expression and purification of BiP, the BiP-PrK-pProEX and pEvol tRNA^Pyl^ PylRS^WT^ plasmids(73) were co-expressed in *E. coli* BL21-AI as described previously (23) with the following modifications: 1 mM N-propargyl-lysine was added after an initial incubation at 37°C for approximately 30 min (OD_600nm_(*E.coli*)=0.2). Induction was achieved by the combined addition of IPTG and 0.02% L-(+)-arabinose.

### Protein labeling

Atto488 dyes were attached to propargyl-lysine by copper-catalyzed Click reactions of 5 nmol BiP-PrK and 25 nmol Atto488-azide in the presence of 2 mM CuSO_4_, 4 mM TBTA, 0.02 mM TCEP, 10 mM fresh sodium ascorbate and protease inhibitor cocktail at room temperature for 2 hours. Centrifugal filters with a cutoff of 10 kDa were used to remove excess dye. The attachment of Atto532-maleimide and Atto647N-maleimide to cysteine residues was performed according to the protocol of the manufacturer (Atto-Tec, Siegen, Germany). Cysteines were reduced by incubation with 1 mM DTT followed by buffer exchange into oxygen-free buffer containing 50 μM TCEP and labeling with two- to threefold molar excess of dye at room temperature for two hours. Unbound fluorophores were removed by a size-dependent filtration. The labeling efficiency was determined via fluorophore absorption and the protein concentration using a bicinchoninic acid assay.

### Protein measurements

Triple labeled BiP was diluted to concentrations of 20-100 pM in 50 mM Hepes, pH7.5, 150 mM KCl, and 10 mM MgCl_2_. Measurements were performed in the absence or in the presence of 1 mM ADP or 1 mM ATP.

## Code availability

Source code for three-color photon distribution analysis is available through the open-source software package *PAM*, freely available under http://www.gitlab.com/PAM-PIE/PAM (71). The module *tcPDA.m* is tightly integrated with PAM, but also supports a text-based file format for direct loading of three-color FRET datasets.

## Acknowledgements

We thank Jelle Hendrix for helpful discussions and Johannes Meyer zum Alten Borgloh for help with implementing computational routines on GPUs. Our thanks belong to Edward Lemke, Swati Tyagi, and Christine Koehler for the introduction to protein expression using non-natural amino acids. We thank Mathias Rosam and Johannes Buchner for providing the BiP plasmid. L.V. acknowledges funding by P-CUBE: The research leading to these results has received funding from the European Community’s Seventh Framework Programme (FP7/2007-2013) under grant agreement n° 227764 (P-CUBE). We gratefully acknowledge the financial support by the Deutsche Forschungsgemeinschaft via the Collaborative Research Consortium on Conformational Switches (SFB1035, Project A11), and by the Ludwig-Maximilians-Universität through the Center for NanoScience (CeNS) and the BioImaging Network (BIN).

## Author contributions

A.B. built the experimental setup, developed and programmed the analysis method, and performed the analysis and measurements of DNA samples. L.V. expressed and labeled proteins and performed the protein measurements. A.B. and D.C.L. wrote the manuscript.

## References

1. Ha T, et al. (1996) Probing the interaction between two single molecules: fluorescence resonance energy transfer between a single donor and a single acceptor. Proc Natl Acad Sci USA 93(13):6264–6268.

2. Myong S, Rasnik I, Joo C, Lohman TM, Ha T (2005) Repetitive shuttling of a motor protein on DNA. Nature 437(7063):1321–1325.

3. Ha T (2001) Single-Molecule Fluorescence Resonance Energy Transfer. Methods 25(1):78–86.

4. Schluesche P, Stelzer G, Piaia E, Lamb DC, Meisterernst M (2007) NC2 mobilizes TBP on core promoter TATA boxes. Nature Structural & Molecular Biology 14(12):1196–1201.

5. Weiss S (1999) Fluorescence spectroscopy of single biomolecules. Science 283(5408):1676–1683.

6. Zander C, et al. (1996) Detection and characterization of single molecules in aqueous solution. Applied Physics B: Lasers and Optics 63(5):517–523.

7. Brooks Shera E, Seitzinger NK, Davis LM, Keller RA, Soper SA (1990) Detection of single fluorescent molecules. Chemical Physics Letters 174(6):553–557.

8. Eggeling C, et al. (2001) Data registration and selective single-molecule analysis using multi-parameter fluorescence detection. Journal of Biotechnology 86(3):163–180.

9. Widengren J, et al. (2006) Single-Molecule Detection and Identification of Multiple Species by Multiparameter Fluorescence Detection. Anal Chem 78(6):2039–2050.

10. Tellinghuisen J, Goodwin PM, Ambrose WP, Martin JC, Keller RA (1994) Analysis of fluorescence lifetime data for single Rhodamine molecules in flowing sample streams. Anal Chem 66(1):64–72.

11. Eggeling C, Fries JR, Brand L, Günther R, Seidel CA (1998) Monitoring conformational dynamics of a single molecule by selective fluorescence spectroscopy. Proc Natl Acad Sci USA 95(4):1556–1561.

12. Fries JR, Brand L, Eggeling C, Köllner M (1998) Quantitative identification of different single molecules by selective time-resolved confocal fluorescence spectroscopy. The Journal of ….

13. Schaffer J, et al. (1999) Identification of Single Molecules in Aqueous Solution by Time-Resolved Fluorescence Anisotropy. J Phys Chem A 103(3):331–336.

14. Kapanidis AN, et al. (2005) Alternating-Laser Excitation of Single Molecules. Acc Chem Res 38(7):523–533.

15. Müller BK, Zaychikov E, Bräuchle C, Lamb DC (2005) Pulsed interleaved excitation. Biophys J 89(5):3508–3522.

16. Hellenkamp B, et al. (2017) Precision and accuracy of single-molecule FRET measurements - a worldwide benchmark study. doi:arXiv:1710.03807v2 [q-bio.QM].

17. Kudryavtsev V, et al. (2012) Combining MFD and PIE for accurate single-pair Förster resonance energy transfer measurements. ChemPhysChem 13(4):1060–1078.

18. Antonik M, Felekyan S, Gaiduk A, Seidel CAM (2006) Separating Structural Heterogeneities from Stochastic Variations in Fluorescence Resonance Energy Transfer Distributions via Photon Distribution Analysis. J Phys Chem B 110(13):6970–6978.

19. Nir E, et al. (2006) Shot-noise limited single-molecule FRET histograms: comparison between theory and experiments. J Phys Chem B 110(44):22103–22124.

20. Kalinin S, Valeri A, Antonik M, Felekyan S, Seidel CAM (2010) Detection of structural dynamics by FRET: a photon distribution and fluorescence lifetime analysis of systems with multiple states. J Phys Chem B 114(23):7983–7995.

21. Tsukanov R, Tomov TE, Berger Y, Liber M, Nir E (2013) Conformational Dynamics of DNA Hairpins at Millisecond Resolution Obtained from Analysis of Single-Molecule FRET Histograms. J Phys Chem B 117(50):131126115511008–16109.

22. Rothwell PJ, et al. (2013) dNTP-dependent conformational transitions in the fingers subdomain of Klentaq1 DNA polymerase: insights into the role of the “nucleotide-binding” state. Journal of Biological Chemistry 288(19):13575–13591.

23. Marcinowski M, et al. (2011) Substrate discrimination of the chaperone BiP by autonomous and cochaperone-regulated conformational transitions. Nature Structural & Molecular Biology 18(2):150–158.

24. Gansen A, et al. (2009) Nucleosome disassembly intermediates characterized by single-molecule FRET. Proceedings of the National Academy of Sciences 106(36):15308–15313.

25. Cristovao M, et al. (2012) Single-molecule multiparameter fluorescence spectroscopy reveals directional MutS binding to mismatched bases in DNA. Nucleic Acids Research 40(12):5448–5464.

26. Santoso Y, Torella JP, Kapanidis AN (2010) Characterizing Single-Molecule FRET Dynamics with Probability Distribution Analysis. ChemPhysChem 11(10):2209–2219.

27. Krainer G, et al. (2017) Ultrafast protein folding in membrane-mimetic environments. Journal of Molecular Biology. doi:10.1016/j.jmb.2017.10.031.

28. Hellenkamp B, Wortmann P, Kandzia F, Zacharias M, Hugel T (2016) Multidomain structure and correlated dynamics determined by self-consistent FRET networks. Nat Meth 14(2):174–180.

29. Rosam M, et al. (2018) Bap (Sil1) regulates the molecular chaperone BiP by coupling release of nucleotide and substrate. Nature Structural & Molecular Biology 25(1):90–100.

30. Horsey I, Furey WS, Harrison JG, Osborne MA, Balasubramanian S (2000) Double fluorescence resonance energy transfer to explore multicomponent binding interactions: a case study of DNA mismatches. Chem Commun 0(12):1043–1044.

31. Ramirez-Carrozzi VR, Kerppola TK (2001) Dynamics of Fos-Jun-NFAT1 complexes. Proc Natl Acad Sci USA 98(9):4893–4898.

32. Liu J, Lu Y (2002) FRET study of a trifluorophore-labeled DNAzyme. J Am Chem Soc 124(51):15208–15216.

33. Watrob HM, Pan C-P, Barkley MD (2003) Two-step FRET as a structural tool. J Am Chem Soc 125(24):7336–7343.

34. Haustein E, Jahnz M, Schwille P (2003) Triple FRET: A tool for Studying Long-Range Molecular Interactions. ChemPhysChem 4(7):745–748.

35. Lee NK, et al. (2007) Three-Color Alternating-Laser Excitation of Single Molecules: Monitoring Multiple Interactions and Distances. Biophys J 92(1):303–312.

36. Person B, Stein IH, Steinhauer C, Vogelsang J, Tinnefeld P (2009) Correlated movement and bending of nucleic acid structures visualized by multicolor single-molecule spectroscopy. ChemPhysChem 10(9-10):1455–1460.

37. Hohng S, Joo C, Ha T (2004) Single-Molecule Three-Color FRET. Biophys J 87(2):1328–1337.

38. Lee J, et al. (2010) Single-molecule four-color FRET. Angew Chem Int Ed Engl 49(51):9922–9925.

39. Lee S, Lee J, Hohng S (2010) Single-Molecule Three-Color FRET with Both Negligible Spectral Overlap and Long Observation Time. PLoS ONE 5(8):e12270.

40. Clamme J-P, Deniz AA (2005) Three-Color Single-Molecule Fluorescence Resonance Energy Transfer. ChemPhysChem 6(1):74–77.

41. Stein IH, Steinhauer C, Tinnefeld P (2011) Single-molecule four-color FRET visualizes energy-transfer paths on DNA origami. J Am Chem Soc 133(12):4193–4195.

42. Chung HS, et al. (2017) Oligomerization of the tetramerization domain of p53 probed by two-and three-color single-molecule FRET. Proceedings of the National Academy of Sciences 114(33):E6812–E6821.

43. Milles S, Koehler C, Gambin Y, Deniz AA, Lemke EA (2012) Intramolecular three-colour single pair FRET of intrinsically disordered proteins with increased dynamic range. Mol BioSyst 8(10):2531–2534.

44. Milles S, et al. (2012) Click strategies for single-molecule protein fluorescence. J Am Chem Soc 134(11):5187–5195.

45. Shi X, et al. (2012) Quantitative fluorescence labeling of aldehyde-tagged proteins for single-molecule imaging. Nat Meth 9(5):499–503.

46. Antos JM, et al. (2009) Site-specific N- and C-terminal labeling of a single polypeptide using sortases of different specificity. J Am Chem Soc 131(31):10800–10801.

47. Hastings WK (1970) Monte Carlo sampling methods using Markov chains and their applications. Biometrika 57(1):97–109.

48. Metropolis N, Rosenbluth AW, Rosenbluth MN, Teller AH, Teller E (1953) Equation of State Calculations by Fast Computing Machines. J Chem Phys 21(6):1087.

49. Förster T (1948) Zwischenmolekulare Energiewanderung und Fluoreszenz. Annalen der Physik 437(1-2):55–75.

50. Lee NK, et al. (2005) Accurate FRET Measurements within Single Diffusing Biomolecules Using Alternating-Laser Excitation. Biophys J 88(4):2939–2953.

51. Hauschild T, Jentschel M (2001) Comparison of maximum likelihood estimation and chi-square statistics applied to counting experiments. Nuclear Instruments and Methods in Physics Research Section A: Accelerators, Spectrometers, Detectors and Associated Equipment 457(1-2):384–401.

52. Kalinin S, Sisamakis E, Magennis SW, Felekyan S, Seidel CAM (2010) On the origin of broadening of single-molecule FRET efficiency distributions beyond shot noise limits. J Phys Chem B 114(18):6197–6206.

53. Bayes T (1763) A Letter from the Late Reverend Mr. Thomas Bayes, F. R. S. to John Canton, M. A. and F. R. S. on JSTOR. Philosophical Transactions (1683-1775). doi:10.2307/105732.

54. Schwarz G (1978) Estimating the Dimension of a Model. The Annals of Statistics 6(2):461–464.

55. Kalinin S, Felekyan S, Antonik M, Seidel CAM (2007) Probability distribution analysis of single-molecule fluorescence anisotropy and resonance energy transfer. J Phys Chem B 111(34):10253–10262.

56. Kalinin S, et al. (2012) A toolkit and benchmark study for FRET-restrained high-precision structural modeling. Nat Meth 9(12):1218–1225.

57. Woźniak AK, Schröder GF, Grubmüller H, Seidel CAM, Oesterhelt F (2008) Single-molecule FRET measures bends and kinks in DNA. Proceedings of the National Academy of Sciences 105(47):18337–18342.

58. Höfig H, Gabba M, Poblete S, Kempe D, Fitter J (2014) Inter-dye distance distributions studied by a combination of single-molecule FRET-filtered lifetime measurements and a weighted accessible volume (wAV) algorithm. Molecules 19(12):19269–19291.

59. Stelzl LS, Erlenbach N, Heinz M, Prisner TF, Hummer G (2017) Resolving the Conformational Dynamics of DNA with Ångstrom Resolution by Pulsed Electron–Electron Double Resonance and Molecular Dynamics. J Am Chem Soc 139(34):11674–11677.

60. Vandenberk N, Barth A, Borrenberghs D, Hofkens J, Hendrix J (2018) Evaluation of Blue and Far-Red Dye Pairs in Single-Molecule FRET Experiments. J Phys Chem B. doi:10.1021/acs.jpcb.8b00108.

61. Doose S, Neuweiler H, Sauer M (2009) Fluorescence quenching by photoinduced electron transfer: a reporter for conformational dynamics of macromolecules. ChemPhysChem 10(9-10):1389–1398.

62. Vogelsang J, et al. (2008) A Reducing and Oxidizing System Minimizes Photobleaching and Blinking of Fluorescent Dyes. Angew Chem Int Ed Engl 47(29):5465–5469.

63. Wang Q, Moerner WE (2013) Lifetime and spectrally resolved characterization of the photodynamics of single fluorophores in solution using the anti-Brownian electrokinetic trap. J Phys Chem B 117(16):4641–4648.

64. Chung HS, Louis JM, Eaton WA (2009) Experimental determination of upper bound for transition path times in protein folding from single-molecule photon-by-photon trajectories. Proceedings of the National Academy of Sciences 106(29):11837–11844.

65. Chung HS, Louis JM, Eaton WA (2010) Distinguishing between protein dynamics and dye photophysics in single-molecule FRET experiments. Biophys J 98(4):696–706.

66. Kityk R, Vogel M, Schlecht R, Bukau B, Mayer MP (2015) Pathways of allosteric regulation in Hsp70 chaperones. Nature Communications 6:8308.

67. Sikor M, Mapa K, Voithenberg von LV, Mokranjac D, Lamb DC (2013) Real-time observation of the conformational dynamics of mitochondrial Hsp70 by spFRET. The EMBO Journal 32(11):1639–1649.

68. Kityk R, Kopp J, Sinning I, Mayer MP (2012) Structure and dynamics of the ATP-bound open conformation of Hsp70 chaperones. Molecular Cell 48(6):863–874.

69. Kapanidis AN, et al. (2004) Fluorescence-aided molecule sorting: analysis of structure and interactions by alternating-laser excitation of single molecules. Proc Natl Acad Sci USA 101(24):8936–8941.

70. Cordes T, Vogelsang J, Tinnefeld P (2009) On the mechanism of Trolox as antiblinking and antibleaching reagent. J Am Chem Soc 131(14):5018–5019.

71. Schrimpf W, Barth A, Hendrix J, Lamb DC (2018) PAM: A Framework for Integrated Analysis of Imaging, Single-Molecule, and Ensemble Fluorescence Data. Biophys J. doi:10.1016/j.bpj.2018.02.035.

72. Tomov TE, et al. (2012) Disentangling Subpopulations in Single-Molecule FRET and ALEX Experiments with Photon Distribution Analysis. Biophys J 102(5):1163–1173.

73. Chatterjee A, Sun SB, Furman JL, Xiao H, Schultz PG (2013) A versatile platform for single-and multiple-unnatural amino acid mutagenesis in Escherichia coli. Biochemistry 52(10):1828–1837.

